# Germ granule higher-order organization coordinates their different functions

**DOI:** 10.1101/2023.11.24.568558

**Authors:** Anne Ramat, Ali Haidar, Céline Garret, Martine Simonelig

**Affiliations:** Institute of Human Genetics, Université de Montpellier, CNRS, Montpellier, France

**Keywords:** Biomolecular condensates, Germ granule, mRNA storage, RNA granule, Suntag, Translation

## Abstract

Most RNP condensates are composed of heterogeneous immiscible phases. However, how this multiphase organization contributes to their biological functions remains largely unexplored. *Drosophila* germ granules, a class of RNP condensates, are the site of mRNA storage and translational activation. Here, using super-resolution microscopy and single-molecule imaging approaches, we show that germ granules have a biphasic organization and that translation occurs in the outer phase and at the surface of the granules. The localization, directionality and compaction of mRNAs within the granule depend on their translation status, translated mRNAs being enriched in the outer phase with their 5’end oriented towards the surface. Altering germ granule biphasic organization represses translation. These findings demonstrate the importance of RNA granule architecture in organizing different functions, highlighting the functional compartmentalization of RNA granules and the key role of higher-order organization.

## Introduction

Membraneless biomolecular condensates have recently emerged as fundamental in cell biology favoring the coordination and efficiency of biochemical reactions by concentrating substrates and enzymes in a common space. Their assembly depends on a demixing process that involves multivalent interactions between proteins and nucleic acids once their concentrations reach a specific threshold ^1^. RNA-protein (RNP) condensates, also called RNA granules, exist in both the nucleus and cytoplasm and are linked to most aspects of RNA biology ^2,3^. Yet, although the biophysical properties and assembly mechanisms of RNP condensates have been deeply investigated, the biological functions of most of them remain poorly understood ^4–6^. Super-resolution microscopy approaches have revealed that many RNP condensates are not homogeneous but rather composed of multiple immiscible phases, reflected by a heterogeneous distribution of RNA and/or RNA binding proteins ^3,7^. The best understood example of this phenomenon is the nucleolus in which three nested phases are linked to three different steps of ribosome biogenesis ^8–10^. For other RNP condensates, the functional relevance of their higher-order organization remains elusive.

Germ granules are evolutionary conserved germline specific RNP condensates that instruct germ cell fate through mRNA regulation ^11,12^. They are hubs for mRNA storage and translational control. *Drosophila* embryonic germ granules contain ≈200 different maternal mRNAs ^13,14^ as well as four main proteins: the germ granule inducer Oskar (Osk), Vasa, an RNA helicase homolog of human DDX4, the scaffold protein Tudor (Tud) and Aubergine (Aub), an Argonaute protein of the PIWI clade ^15^. *Drosophila* germ granules display both liquid-like and hydrogel-like properties, mRNA representing a stable component of the granule ^16,17^. Multiple copies of the same mRNA molecules are grouped together in homotypic clusters ^18,19^, although the function behind this organization is still unknown. In several species, germ granules have a multiphase architecture ^20,21^, however, the relationships between this organization and germ granule functions have not been addressed. Furthermore, although the function of germ granules in the storage of repressed mRNAs is well described, whether they are also sites for translational activation remains an open question. In *C. elegans*, germ granule mRNA translation takes place after their exit from the granule and dispersion into the cytoplasm ^22,23^. Strikingly, in *Drosophila* embryos, translation of germ granule mRNAs follows a tight temporal sequence ^13^, implying that germ granules achieve temporal translational control by coordinating concomitant translational repression of some mRNAs and activation of others. We took advantage of this outstanding property to determine how these opposite functions are compartmentalized within germ granules, and address the relationships between architecture and biological functions of RNP condensates.

## Results

### *Drosophila* germ granules have a biphasic organization

Due to the small size of germ granules, ranging from 200 to 500 nm ^24^, we used super-resolution 2D Stimulated Emission Depletion (STED) microscopy to characterize the distribution of germ granule main protein components. We set up a method to slice and expose the posterior pole of embryos to the objective, in order to reduce the distance between the sample and the objective and get optimal resolution (Fig. S1A-C and Supplementary Text). All four main components of germ granules were enriched in the outer most part of the granule that we termed outer phase or shell (Fig. 1A) and they colocalized as quantified using the Pearson Correlation Coefficient (PCC(Costes)) (Fig. S1D). This organization was reminiscent of the germ granule structure observed previously by electron microscopy, showing an electron dense periphery and lucid center^25,26^. STED microscopy requires the use of an antibody to get a signal that stands the depletion. To rule out that the observed pattern was the consequence of poor antibody penetration within the granule, we used AiryScan and 3D-OMX microscopy to visualize the localization of GFP-tagged versions of Aub, Vasa and Tud using GFP fluorescence. The same enrichment was observed in the shell (Fig. 1B, Fig. S1C). Furthermore, GFP-Aub and Vasa-tdTomato visualized using 3D-OMX without antibody staining also colocalized in the shell of germ granules (Fig. 1C). GFP-Aub enrichment in the outer phase was visible in 3D (Fig. S1E), consistent with previous observations based on electron microscopy of ultra-thin sections that led to propose a hollow sphere morphology of germ granules ^26^. Using STED acquisitions and Osk protein as a marker, we measured the size of germ granules and their outer phase. Germ granule diameters ranged from 110 to 600 nm with an average of 315 nm (Fig. 1D), and the thickness of the outer phase ranged from 12 to 279 nm with an average of 112 nm (Fig. 1E). We conclude that germ granules have a multiphase structure, with main protein components enriched in the outer phase.

**Fig. 1.**
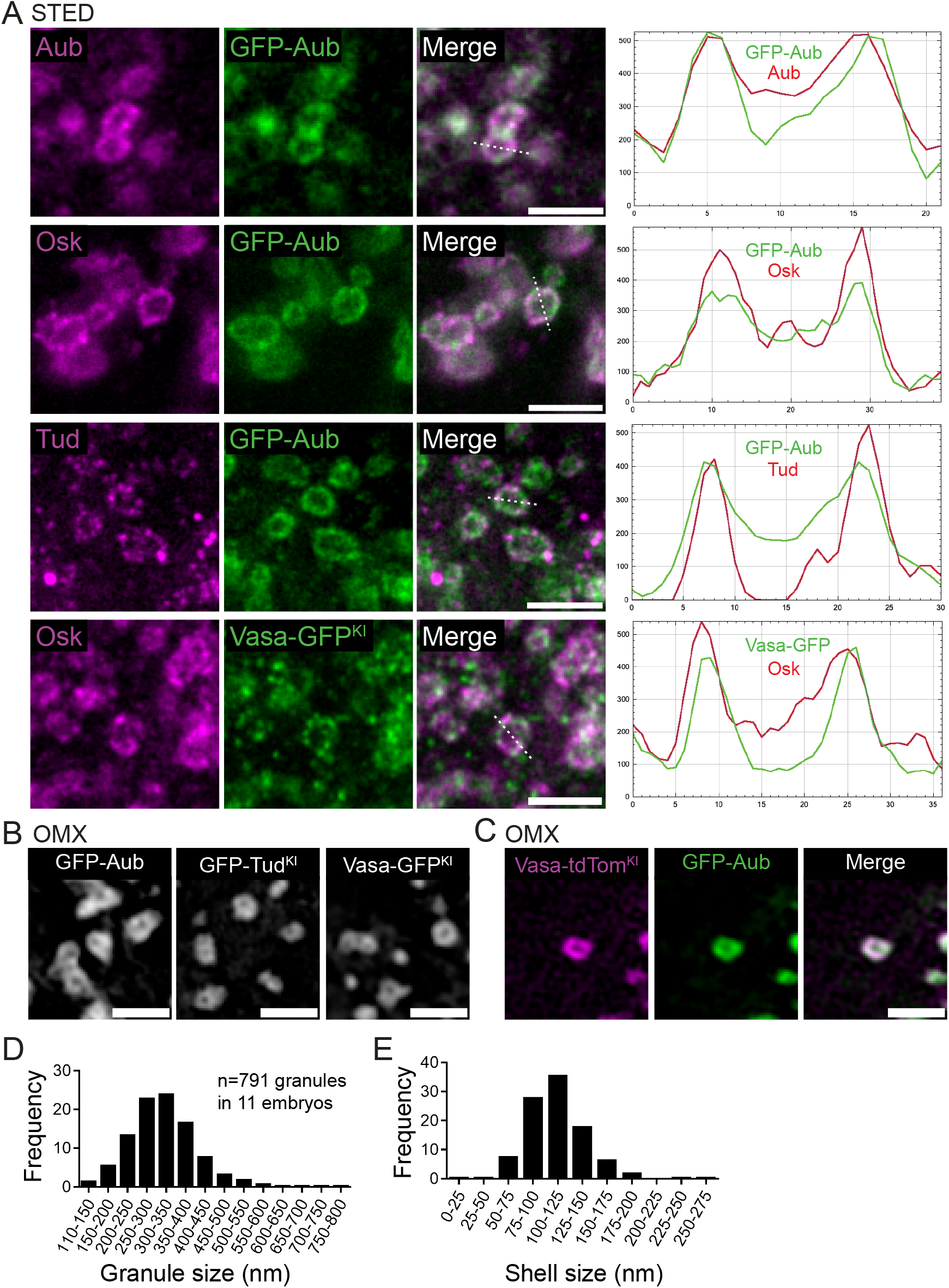
*Drosophila* germ granules have a biphasic organization. (**A**) STED imaging of germ granule main components. Immunostaining of *UASp-GFP-Aub; nos-Gal4* or *vasa-GFP^KI^* embryos with anti-Aub (magenta), anti-Osk (magenta), anti-Tud (magenta) and anti-GFP (green) to visualize GFP-Aub or Vasa-GFP. Fluorescence intensity (right) was measured along the path marked with a white dashed line and intensity profiles of each channel as indicated on the graph. (**B**) 3D-OMX imaging of *UASp-GFP-Aub; nos-Gal4* (left), *GFP-Tud^KI^*(middle), *Vasa-GFP^KI^* (right) in which GFP was directly recorded without antibody staining. (**C**) 3D-OMX imaging of *UASp-GFP-Aub/ vasa-tdTom^KI^; nos-Gal4/+* embryos in which GFP and tdTomato were directly recorded without antibody staining. (**D-E**) Measurement of germ granule size (**D**) and outer phase size (**E**) from wild-type embryos immunostained with Osk antibody and imaged using STED microscopy. Scale bars: 1µm.

### Aub and mRNAs are required for the biphasic architecture and maintenance of germ granules

Aub binds germ granule mRNAs through base-pairing with piRNAs and is a major actor of their recruitment to germ granules ^14,27,28^. Through this function, Aub is expected to play an important role in maintaining the RNA content of germ granules. We thus addressed the contribution of mRNAs to germ granule biphasic organization by analyzing germ granule architecture in the absence of Aub. We used a specific genetic set up to visualize germ granules in *aub* mutant embryos. Indeed, due to early developmental defects during oogenesis in *aub* mutants, Osk protein is not produced at the posterior pole, and germ granules do not form in these mutants ^29^. However, Osk does accumulate at the anterior pole of embryos following recruitment of *osk* mRNA to the anterior pole, even in the absence of Aub ^27,30^. In this context, germ granule mRNAs localize very poorly to the anterior pole, consistent with the role of Aub in mRNA recruitment to germ granules ^27^. Using STED imaging and Osk as a marker, we found that germ granules produced at the anterior pole when *osk* mRNA was delocalized had a similar biphasic organization as those present at the posterior pole of wild-type embryos (Fig. S2A). In contrast, in *aub* mutant embryos, germ granule architecture was severely affected, with the disappearance of their biphasic organization and a diameter reduced to 270 +/− 43 nm, instead of 387 +/− 63 nm in the *aub^+^* background (Fig. S2B, C). In lines with Aub function in mRNA recruitment to germ granules, *nos* mRNA was not found in these germ granules using smFISH and STED imaging (Fig. S2D). In addition, these germ granules disappeared altogether in 79.3% of embryos (n=53) after 2.5 hours of development, while they were maintained in the *aub^+^* background (100% of embryos, n=50).

These data demonstrate the essential contribution of Aub and mRNAs to germ granule biphasic organization and to their maintenance.

### *nos* mRNA translation occurs in the outer phase, and at the surface and immediate periphery of germ granules

To understand how germ granule mRNA translation integrates within germ granule architecture, we used the Suntag method that allows the visualization of ongoing translation ^31–35^, to monitor *nos* mRNA translation. Briefly, the binding of single-chain antibodies fused to GFP (scFv-GFP) on an array of GCN4-based epitope (called Suntag) placed downstream of the start codon of the mRNA of interest allows the visualization of nascent peptides by creating bright GFP spots above background whose intensity reflects translation efficiency (Fig. 2A). Translating mRNAs are detected either by the MS2/MCP system ^36^ or single molecule Fluorescent In Situ Hybridization (smFISH) ^37^. We inserted an array of 12 Suntag repeats after *nos* initiation codon (Fig. S3A) in a genomic construct known to rescue *nos* RNA null mutant (*nos^BN^*) phenotypes ^38^. We first recorded the efficiency of *suntag-nos* mRNA localization to the germ plasm, the cytoplasm containing germ granules at the posterior pole of embryos, by measuring *suntag* smFISH fluorescence intensity at the posterior pole and in whole embryos expressing *suntag-nos*, and comparing this quantification to that of *nos* smFISH fluorescence intensity in wild-type embryos. While 3.8% of *nos* mRNA localized to the posterior pole in wild-type embryos as previously reported ^18,39^, 2.15% of *suntag-nos* mRNA localized to the posterior pole (Fig. S3B), indicating a reduced recruitment of the chimeric *suntag-nos* mRNA to the germ plasm. Nonetheless, the *suntag-nos* construct was able to rescue *nos^BN^* embryonic lethality (*nos^BN^*: 100% embryonic lethality, *suntag-nos/+; nos^BN^*: 48.7% embryonic lethality, n>100 embryos), showing that the regulation of *suntag-nos* mRNA, including its level of localization and translational control allowed embryonic development.

**Fig. 2.**
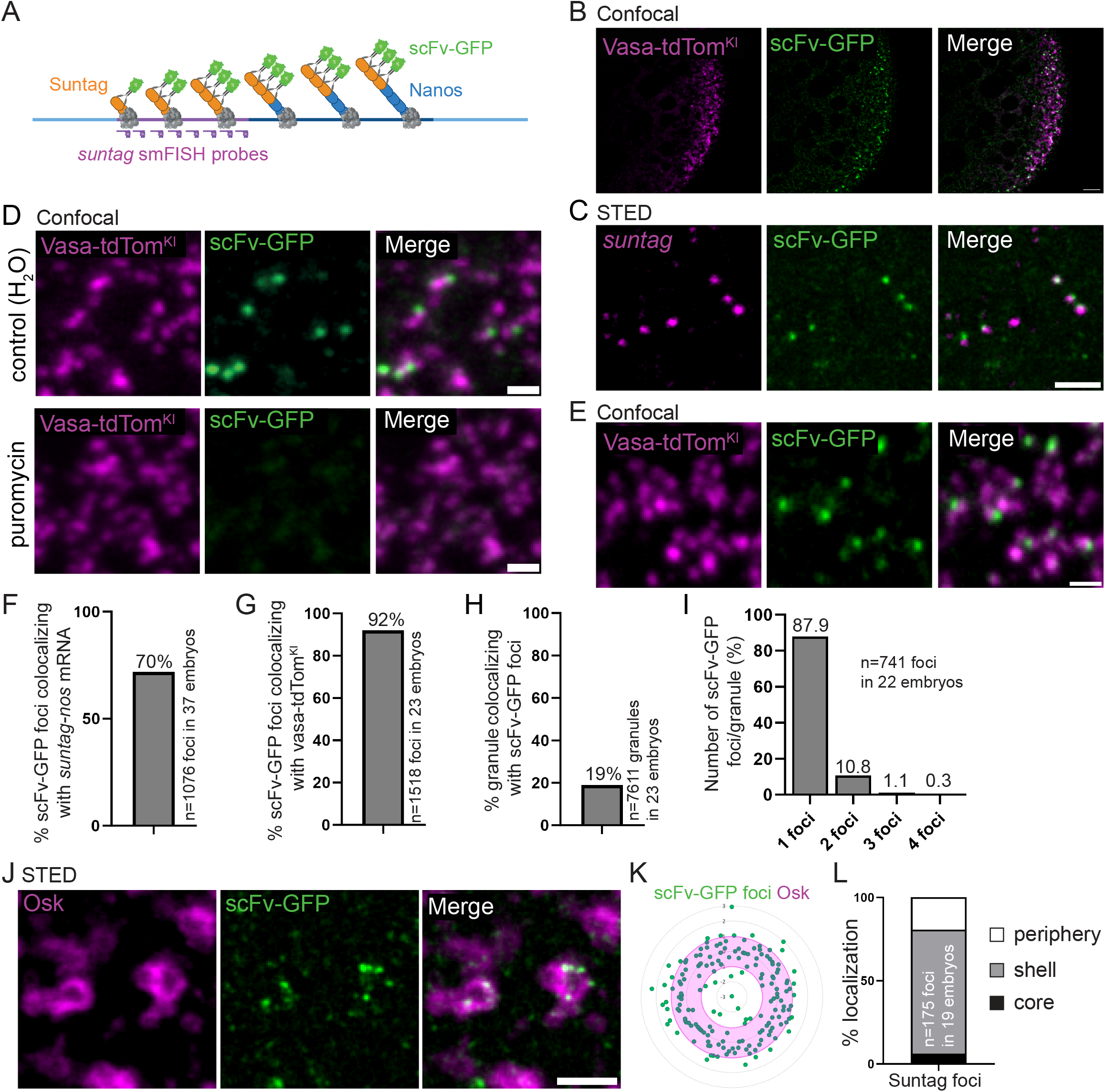
Visualization of *nos* mRNA translation at germ granules using Suntag. (**A**) Principle of the Suntag technique to visualize *nos* mRNA translation. Blue lines represent *nos* 5’ and 3’UTRs, the purple line represents the Suntag array, and the dark blue line represents *nos* coding sequence. The polysomes are shown in grey. The Suntag and Nos nascent peptides are shown as orange and blue circles, respectively. The scFv-GFP antibody in green binds the Suntag peptide. *suntag* smFISH probes are shown in purple. (**B**) Confocal images of *vasa-tdTom^KI^/suntag-nos; scFv-GFP/+* embryos. Vasa-tdTomato (magenta) and scFv-GFP (green) fluorescence were recorded directly without antibody staining. (**C**) STED images of immuno-smFISH of *suntag-nos/+; scFv-GFP/+* embryos with anti-GFP nanobody (green) to detect scFv-GFP and *suntag* smFISH probes (magenta). (**D**) Fluorescent confocal images of *vasa-tdTom^KI^/suntag-nos; scFv-GFP/+* permeabilized embryos without drug treatment (top) and with puromycin treatment (bottom). Vasa-tdTomato (magenta) and scFv-GFP (green) fluorescence were recorded without antibody staining. (**E**) Close-up view of fluorescent confocal images of *vasa-tdTom^KI^/suntag-nos; scFv-GFP/+* embryos showing scFv foci with germ granules. Vasa-tdTomato (magenta) and scFv-GFP (green) fluorescence were recorded without antibody staining. (**F**) Percentage of scFv-GFP foci colocalizing with *suntag-nos* mRNA obtained from images as in (**C**). (**G**) Percentage of scFv-GFP foci colocalizing with germ granules marked with Vasa-tdTom obtained from images as in (**E**). (**H**) Percentage of germ granules marked with Vasa-tdTom colocalizing with scFv-GFP foci, i.e. undergoing translation, obtained from images as in (**E**). (**I**) Quantification of the number of scFv-GFP foci per granule obtained from images as in (**E**). (**J**) STED imaging of *suntag-nos/+; scFv-GFP/+* embryos immunostained with anti-Osk antibody (magenta) and anti-GFP nanobody (green) to reveal scFv-GFP. (**K**, **L**) Quantification of svFv-GFP foci localization within the germ granule biphasic structure, obtained from STED images as in (**J**). Radar plot of the relative distance of scFv-GFP foci (green dots) within Osk immunostaining (magenta). The hollow circle represents the germ granule outer phase (**K**). Percentage of scFv-GFP foci localized in the core (black), in and at the surface of the shell (grey) and at the immediate granule periphery (white). Scale bars: 5 µm in (**B**), 1 µm in (**C-E**, **J**).

Using confocal microscopy, we found that *suntag-nos* embryos containing maternally-provided scFv-GFP (*nos-scFv-GFP* ^40^) showed bright GFP foci at the posterior pole as expected for Nos protein production site (Fig. 2B). To determine whether these GFP foci reflected *suntag-nos* mRNA translation, first we analyzed their colocalization with cognate mRNA using smFISH probes directed against the *suntag* sequence, and STED super-resolution imaging (Fig. 2C). We quantified that 70% of scFv-GFP foci colocalized with *suntag* mRNA (Fig. 2F), revealing translating mRNAs. Second, we showed that these scFv-GFP foci disappeared upon treatment of embryos with puromycin that releases translating ribosomes ^31^ (Fig. 2D), confirming that scFV-GFP foci corresponded to translation sites or accumulation of freshly produced Suntag-Nos protein. We next addressed the localization of scFv-GFP foci in relation with germ granules marked with Vasa-tdTomato. Confocal imaging revealed that 92% of scFv-GFP foci colocalized with or overlapped germ granules (Fig. 2E, G). This suggested that *suntag-nos* mRNA translation occurred exclusively at germ granules and not in the intergranular space. To confirm this result, we performed immuno-smFISH to visualize *suntag-nos* mRNA, scFv-GFP foci and germ granules marked with Osk (Fig. S3C). Quantification of confocal images (see Methods) showed that all (99.85%) translation events were restricted to the germ plasm and adjacent or colocalizing with germ granules (Fig. S3C). Therefore *suntag-nos* mRNA translation occurred specifically at germ granules.

We quantified that 19% of germ granules marked with Vasa-tdTomato colocalized with scFv-GFP foci, and were thus translating *suntag-nos* mRNA (Fig. 2 E, H). Most translating germ granules (87.9%) had a single scFv-GFP foci, but up to four scFv-GFP foci could be visualized per granule (Fig. 2I). However, it should be noted that these numbers would be expected to be higher for endogenous *nos* mRNA translation as a lower level of *suntag-nos* localized to the germ plasm (Fig. S3B), and we quantified using FISH-quant ^41^ a lower number of *suntag-nos* mRNA molecules per cluster compared to that of endogenous *nos* mRNA ^42^ (Fig. S3D).

We next used STED super-resolution microscopy to record the localization of translation events within germ granules (Fig. 2J). Strikingly, using a Laplacian of Gaussian filter to define the edges of germ granule outer and inner phases (see Methods), we found that most scFv-GFP foci localized in the outer phase and at the granule immediate periphery (74.3% and 19.4%, respectively), with only 6.3% localizing in the inner phase of the granule (Fig. 2J-L). We performed immuno-smFISH to visualize using STED microscopy, scFv-GFP, *suntag-nos* mRNA and germ granule marked by Osk. We observed that *suntag-nos* mRNA and scFv-GFP translation foci were not organized in a specific order within the outer phase, scFv-GFP foci appearing either at the same level, or in a more internal or more external position than *suntag-nos* mRNA within the outer phase (Fig. S3E). Importantly, when *suntag-nos* mRNA was detected in the inner phase or core of the granule, it was not associated with scFv-GFP foci (Fig. S3F, 100% n=31 granules), indicating a lack of translation.

We conclude that mRNA translation is compartmentalized within germ granules, occurring in the outer phase and at the surface and immediate periphery of the granules. In addition, mRNAs localized in the core of the granules are translationally repressed.

### Translational repressors are not present within germ granules

The regionalization of *suntag-nos* mRNA translation within germ granules implied that translational regulators should be heterogeneously distributed within germ granules. We previously reported a role of Aub in *nos* mRNA translation activation through interaction with the translation initiation factors eIF3d and PABP that localized at the edge of germ granules ^43^. Using STED imaging, we found that eIF3d and PABP accumulated in the outer phase and at the surface of germ granules, but not in the inner phase (Fig. S4A, B), and they colocalized with GFP-Aub using PCC(Costes) (Fig. S4J). We next investigated the localization of translational repressors known to regulate *nos* mRNA in early embryos. Whereas *nos* mRNA translation is activated in the germ plasm, it is repressed in the somatic part of early embryos through deadenylation by the CCR4-NOT deadenylation complex, and by a repressor complex containing Smaug, Cup, Trailer Hitch, and the RNA helicases Me31B -a DDX6 homolog- and Belle -a DDX3 homolog-^44^. We analyzed the localization of Smaug, Cup, Me31B and Belle in the germ plasm using STED microscopy and did not find colocalization of any of these proteins with the germ granule markers Osk or Aub using (PCC(Costes) (Fig. S4C, D, E, G, J). In particular, these translational repressors never accumulated in the core of germ granules where repressed mRNAs were present. However, we could observe Smaug, Cup and Me31B foci at the surface germ granules (Fig. S4C, D, E). Quantification of the distance between Smaug foci and germ granules (see Methods), revealed a significant localization of Smaug in close proximity to germ granules, with 28.3% of Smaug foci within 50 nm of germ granules (Fig. S4K). Smaug is known to interact with Osk in the germ plasm, preventing Smaug binding to *nos* mRNA and relieving translational repression ^45,46^. Given Smaug foci localization around germ granules (Fig. S4C, K), the surface of germ granule might be the site where the Smaug-Osk interaction takes place. Me31B was previously reported to be a component of germ granules ^47^. Although Me31B was not found within germ granules in the germ plasm, it accumulated in the outer phase of germ granules at a later stage when germ granules enlarge by fusion in primordial germ cells (Fig. S4F), consistent with a recent study describing Me31B enrichment in germ granules at this stage ^48^. We then addressed the relationships of the deadenylation machinery with germ granules by recording the localization of two subunits of the deadenylation complex, Not1 and CCR4, in the germ plasm. None of the subunits showed association with germ granules marked with Osk or Aub (Fig. S4H-J).

The lack of translational repressors inside germ granules suggests that translational repression of mRNAs accumulating in the core of germ granules might occur in the absence of repressors and could result from high mRNA compaction and partitioning of translational activators into the shell.

### mRNA localization within the germ granule biphasic architecture depends on their translation status

Translation of the germ granule mRNAs, *nos*, *germ cell less* (*gcl*), *polar granule component* (*pgc*) and *cyclin B* (*cycB*) is sequential ^13^: *nos* translation is already active at the onset of embryogenesis; *gcl* translation starts soon after, Gcl protein being visible 20 minutes (min) after embryo deposition at 25°C ^49^; *pgc* translation takes place after primordial germ cell formation, 90 min after embryo deposition ^13^; and *cycB* translation starts after primordial germ cell incorporation into the gonads, at 12 to 13 hours of embryogenesis ^50^. To understand how these mRNAs integrate within the biphasic organization of germ granules in relation with their translation status, we analyzed their localization using immuno-smFISH with anti-Osk immunostaining to visualize germ granules, combined with STED microscopy. We found that *gcl*, *pgc* and *cycB* mRNAs formed one to two foci within germ granules, consistent with their assembly in homotypic clusters ^18,19^, and that *nos* mRNA assembled in multiple foci, as previously reported ^19^, within and at the immediate periphery of germ granules (Fig 3A-D). We used the number of nuclei to carefully stage embryos and classify them into early (≤2 nuclei, age 0-20 min) and late (>2 nuclei and before primordial germ cell formation, age 20-90 min) stages. The localization of mRNA foci within germ granules was recorded in early and late stage embryos, using the Laplacian of Gaussian filter to define the edges of germ granule core and shell (see Methods). Each mRNA focus was categorized as localized in the core, the shell, or at the immediate periphery of germ granules. The localization of *nos* mRNA whose translation was active in both early and late stage embryos did not significantly change between both time points (Fig. 3E, E’). Similarly, the localization of *pgc* and *cycB* mRNAs whose translation started beyond the developmental period analyzed, did not change between both stages (Fig. 3G, G’, H, H’). In contrast, *gcl* mRNA localization changed after the onset of its translation (Fig. 3F, F’). While in early stage embryos, 30% of *gcl* mRNA foci localized in the core of germ granules, this percentage decreased to 13.5% in late stage embryos. Therefore, upon translation, *gcl* mRNA relocalized from the core to the periphery of germ granules, showing that the distribution of mRNAs within germ granule correlated with their translation status.

**Fig. 3.**
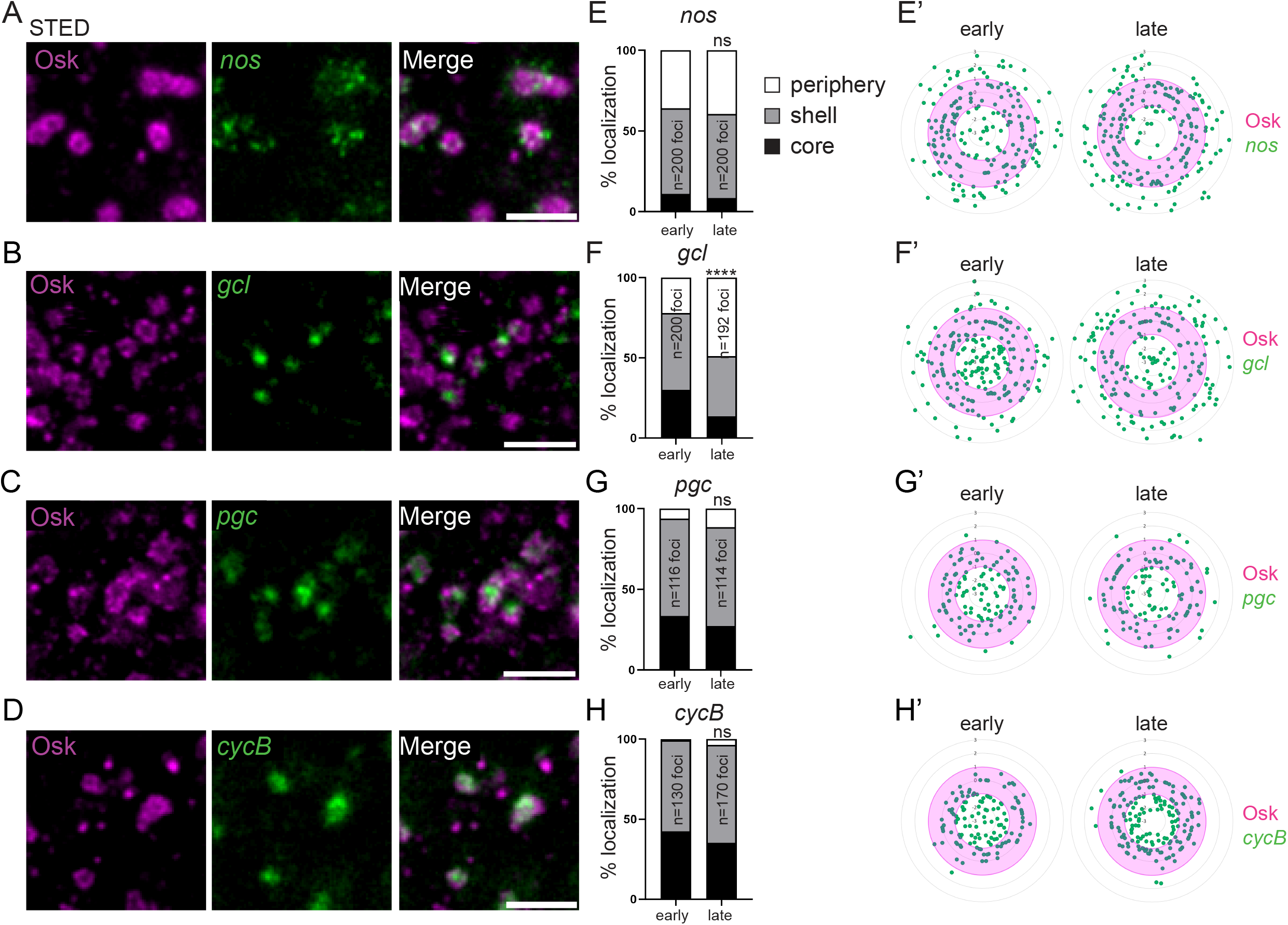
mRNA localization within germ granules. (**A-D**) STED images of immuno-smFISH of wild-type embryos (20-120 min) with anti-Osk (magenta) and smFISH probes (green) against *nos* (**A**), *gcl* (**B**), *pgc* (**C**) and *cycB* (**D**) mRNAs. (**E-H**) Percentage of localization of *nos*, *gcl*, *pgc* and *cycB* mRNA foci in early (0-20 min) and late (20-90 min) wild-type embryos, in the core (black), in and at the surface of the shell (grey) and at the immediate periphery (white) of germ granules from images as in (**A-D**). ns: non-significant, **** *p*<0.0001 using the χ^2^ test. (**E’-H’**) Radar plots of the relative localization of *nos*, *gcl*, *pgc* and *cycB* mRNAs (green dots) within Osk immunostaining (magenta) in early (0-20 min) and late (20-90 min) embryos from images as in (**A-D**). Hollow circles represent the germ granule outer phase. Scale bars: 1 µm.

To address mRNA orientation in relation with their translation within germ granules, we next designed smFISH probes directed against the 5’end and 3’end of *nos* (5’-*nos* and *3’-nos*) as an example of translated mRNA, and *cycB* (5’-*cycB* and 3’-*cycB*) as an example of repressed mRNA (Fig. S5), and performed double smFISH or immuno-smFISH with anti-Osk antibody of embryos. Strikingly, *nos* mRNA 3’ends assembled in a single cluster per granule, surrounded by many smaller foci corresponding to *nos* 5’ends (Fig. 4A). With regards to germ granule core/shell organization defined by Osk staining, *nos* 5’ends were predominantly localized in the shell and at the immediate periphery of germ granules, whereas *nos* 3’ends localized within the core and shell of germ granules (Fig. 4B-E). Intriguingly, *cycB* mRNA showed the reverse orientation. *cycB* 5’ends were localized more internally than their 3’ends (Fig. 4F), and when compared to germ granule organization, *cycB* 5’ends localized within the core and shell of germ granules, whereas its 3’ends localized more externally in the shell and at the periphery of the granules (Fig. 4G-J).

**Fig. 4.**
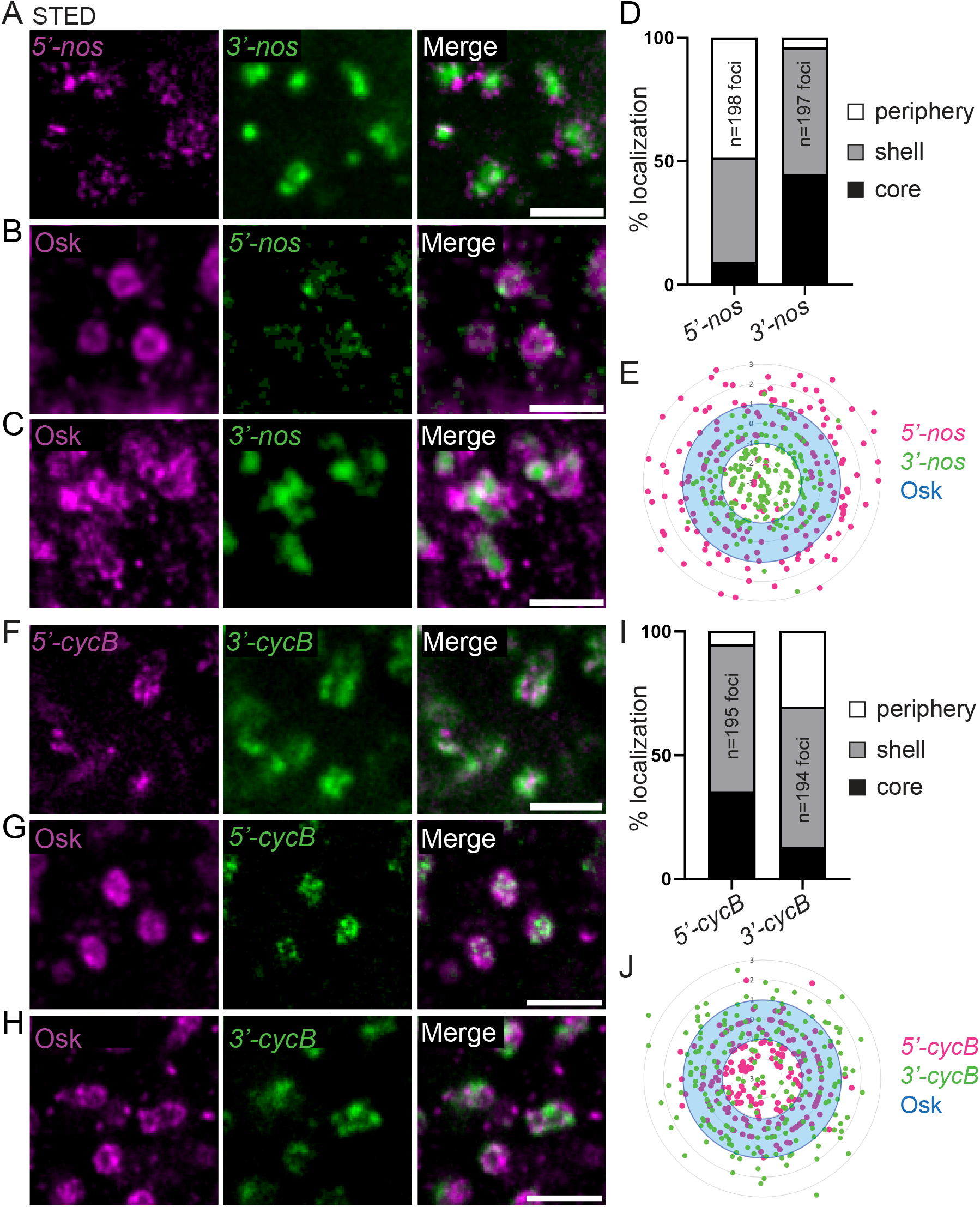
mRNA orientation within germ granules. (**A**) STED images of smFISH of wild-type embryos with *nos* 5’end (5’-*nos*, magenta) and 3’end (3’-*nos*, green) probes. (**B**, **C**) STED images of immuno-smFISH with anti-Osk antibody (magenta) as a marker of germ granules and *nos* 5’end (**B**) or 3’end (**C**) probes (green). (**D**) Percentage of *nos* 5’end and 3’end foci localized in the core (black), in and at the surface of the shell (grey) and at the immediate periphery (white) of germ granules from images as in (**B**, **C**). (**E**) Radar plot of the relative localization of *nos* 5’end (magenta dots) and 3’end (green dots) within Osk immunostaining (blue) in wild-type embryos from images as in (**B**, **C**). The blue hollow circle represents the germ granule outer phase. (**F**) STED images of smFISH of wild-type embryos with *cycB* 5’end (5’-*cycB*, magenta) and 3’end (3’-*cycB*, green) probes. (**G**, **H**) STED images of immuno-smFISH with anti-Osk antibody (magenta) as a marker of germ granules and *cycB* 5’end (**G**) or 3’end (**H**) probes (green). (**I**) Percentage of *cycB* 5’end and 3’end foci localized in the core (black), in and at the surface of the shell (grey) and at the immediate periphery (white) of germ granules from images as in (**G**, **H**). (**J**) Radar plot of the relative localization of *cycB* 5’end (magenta dots) and 3’end (green dots) within Osk immunostaining (blue) in wild-type embryos from images as in (**G**, **H**). The blue hollow circle represents the germ granule outer phase. Scale bars: 1 µm.

Together, these results show mRNAs translocate towards the external regions of germ granules upon translation and that they have a specific orientation within the granules, 5’ends of translated mRNAs being localized externally.

### Reducing translation increases mRNA compaction in germ granules

Egg activation that occurs upon egg laying triggers a massive remodeling of the maternal transcriptome landscape, involving drastic changes in mRNA stability and translation efficiency ^51,52^. The Pan gu (Png) kinase is a major effector of egg activation-mediated posttranscriptional regulation through the phosphorylation of several translational repressors ^53^. We addressed whether *nos* mRNA translation at germ granules would be affected in *png* mutant embryos, making them an excellent genetic system to analyze the links between mRNA translation and localization within germ granules, if it was the case. Using immuno-smFISH and confocal imaging to visualize *suntag-nos* mRNA and scFv-GFP foci, we found a significant decrease in the number of *suntag-nos* mRNA clusters colocalizing with scFv-GFP, i.e. undergoing translation, in *png* mutant compared to control embryos (Fig. S6A, B), whereas the number of *suntag-nos* mRNA molecules per cluster was not reduced (Fig. S6C). Therefore, *suntag-nos* mRNA translation was reduced in *png* mutant embryos. We analyzed the distribution of *nos* and *gcl* mRNAs within germ granules in *png* mutant embryos using immuno-smFISH with anti-Osk antibody and STED imaging. Remarkably, both *nos* and *gcl* mRNA localization was affected in *png* mutant embryos where the percentage of mRNA foci localized in the core of germ granules increased at the expense of that at the periphery (Fig. 5A-D). In contrast, the localization within germ granules of *cycB* mRNA that was not translated at this stage, did not change in *png* mutant embryos (Fig. S7A, B). Thus, decreasing translation led to a redistribution of translating mRNAs towards the inside of germ granules.

**Fig. 5.**
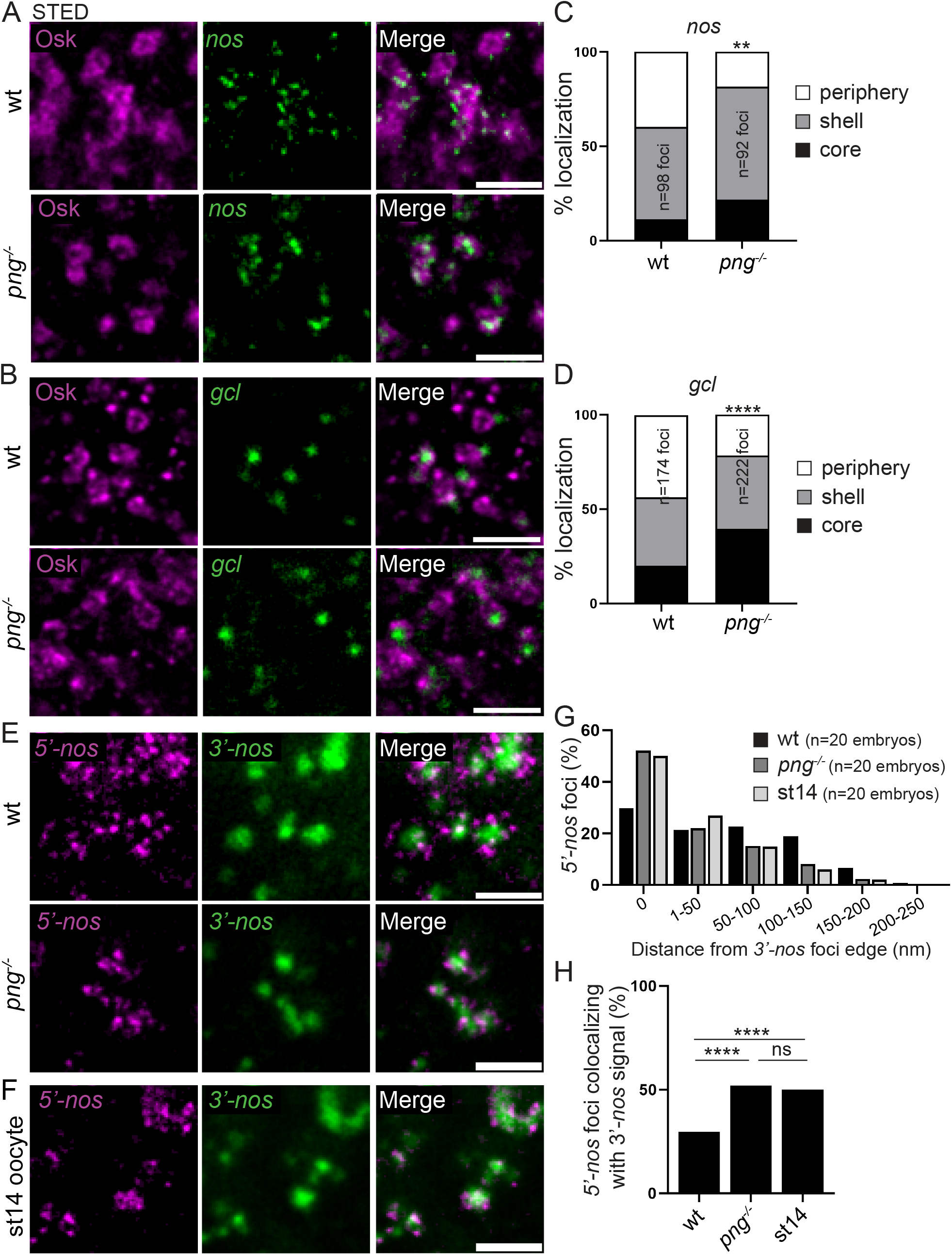
mRNA localization and compaction within germ granules depend on their translation. (**A, B**) STED imaging of immuno-smFISH of wild-type (top) and *png^1058^* (*png^−/−^*, bottom) embryos with anti-Osk antibody (magenta) as a marker of germ granule and smFISH probes (green) against *nos* (**A**) and *gcl* (**B**) mRNAs. (**C**, **D**) Percentage of localization of *nos* (**C**) and *gcl* (**D**) mRNA foci in wild-type and *png^1058^* mutant embryos, in the core (black), in and at the surface of the shell (grey) and at the immediate periphery (white) of germ granules from images as in (**A**, **B**). ** *p*<0.01, **** *p*<0.0001 using the χ^2^ test. (**E**) STED images of smFISH against *nos* 5’end (5’-*nos*, magenta) and 3’end (3’-*nos*, green) in wild-type and *png^1058^* mutant embryos. (**F**) STED images of smFISH against *nos* 5’end (5’-*nos*, magenta) and 3’end (3’-*nos*, green) in wild-type stage 14 oocytes. (**G**) Measurement of the distance between *nos* 5’end foci and the edge of *nos* 3’end foci in wild-type embryos, *png^1058^* embryos and wild-type stage 14 oocytes (st14), from images as in (**E**, **F**). The histogram shows the percentage of *nos* 5’end foci in each distance class. (**H**) Percentage of *nos* 5’end foci colocalizing with *nos* 3’end foci in wild-type embryos, *png^1058^* embryos and wild-type stage 14 oocytes (st14). ns: non-significant, **** *p*<0.0001 using the χ^2^ test. Scale bars: 1 µm.

We next analyzed the orientation and conformation of *nos* mRNA within germ granules upon reduced translation using double smFISH with *nos* 5’end and 3’end specific probes and STED imaging. The 5’-3’ orientation of *nos* mRNA within germ granules did not change in *png* mutant embryos, *nos* 5’end remaining localized externally to *nos* 3’end (Fig. 5E). However, both ends appeared closer to each other than in wild-type embryos. We measured the distance between *nos* 5’end signal and the edge of *nos* 3’end signal (see Methods and Fig. S8) and observed that the proportion of *nos* 5’end colocalizing with *nos* 3’end (distance=0 nm) increased in *png* mutant compared to wild-type embryos (Fig. 5G, H), at the expense of *nos* 5’end separated from *nos* 3’end by more than 50 nm. A similar increase in mRNA compaction was not observed with *cycB* mRNA in *pgn* mutant embryos (Fig. S7C-E). Furthermore, the distance between 5’end and 3’end for *cycB* was closer than that for *nos* mRNA in wild-type embryos, and in the same range as that for *nos* mRNA in *png* mutant embryos (compare Fig. 5G, H and Fig. S7D, E).

In addition to *png* mutant embryos, we took advantage of another context where translation is reduced. A previous report based on GFP-Nos fusion protein reported that *nos* mRNA translation started at the posterior pole during oogenesis, in stage 13 oocytes ^54^. However, we found recording ongoing translation of *suntag-nos* mRNA that it did not occur in stage 14 oocytes (Fig. S9), suggesting that *nos* translation at germ granules was initiated or strongly increased following egg activation. We found that the distance between *nos* 5’end and 3’end was closer in stage 14 oocytes than in embryos, and similar to that in *png* mutant embryos (Fig. 5F-H).

These results strengthen the conclusion that mRNAs move towards the outer phase and periphery of germ granules when they are translated. They also reveal that mRNA conformation depends on their translational status, showing higher compaction when they are not translated and decompaction upon translation.

### Germ granule biphasic organization promotes mRNA translation

We sought to perturb germ granule architecture in order to address its contribution to germ granule functions. We took advantage of a single point mutant of *tud* in the first Tudor domain, *tud^A36^* (Fig. S10A), in which germ granules were described as rods instead of hollow spheres by electron microscopy ^26^. Tud protein contains 11 Tudor domains. These domains are known to mediate interactions with other proteins, in particular through dimethylated arginines. Tudor domains in Tud have been proposed to serve as docking platforms for germ granule assembly ^26^, and indeed the multiple Tudor domains should generate multivalent interactions known to be instrumental in the formation of condensates by phase separation ^55^. Aub undergoes symmetric arginine dimethylation and Tud recruits Aub to germ granules through interaction between several Tudor domains and Aub dimethylated arginines ^56–59^. Interestingly, in *tud^A36^* mutant embryos, although germ granule mRNAs were shown to localize to the posterior pole, the formation of germ cells was strongly affected ^26^, suggesting a defect in mRNA translation at germ granules. Analysis of germ granules in *tud^A36^* embryos, using Osk immunostaining and STED imaging revealed a drastic alteration of germ granule organization. The biphasic structure was lost (Fig. 6A) and the size of germ granule was reduced to an average of 203 +/− 42 nm instead of 313 +/−55 nm in wild-type embryos (Fig. 6B). Investigating *suntag-nos* translation in *tud^A36^* mutant embryos using immuno-smFISH and confocal imaging to record *suntag-nos* mRNA clusters and scFv-GFP foci, we found a strong decrease in translation (Fig. 6C). The percentage of translating *suntag-nos* mRNA clusters, i.e. colocalizing with scFv-GFP foci, decreased from 24.4% in wild-type embryos to 8.5% in *tud^A36^* mutant embryos (Fig. 6D), whereas *suntag-nos* mRNA localization to germ granules was only slightly reduced (mean of 2.4 *suntag-nos* molecules per cluster in wild-type germ granules versus 2.1 in *tud^A36^*) (Fig. S10B). We analyzed the localization of *suntag-nos* translation within germ granules using immunostaining to reveal scFv-GFP foci and Osk as a germ granule marker, visualized with STED microscopy. We confirmed a lower number of scFv-GFP foci in germ granules in *tud^A36^* embryos and these foci localized mostly at the periphery of germ granules or close to their surface (Fig. 6E, Fig. S10C), indicating that the structure of *tud^A36^* germ granules was not compatible with translation inside the granules.

**Fig. 6.**
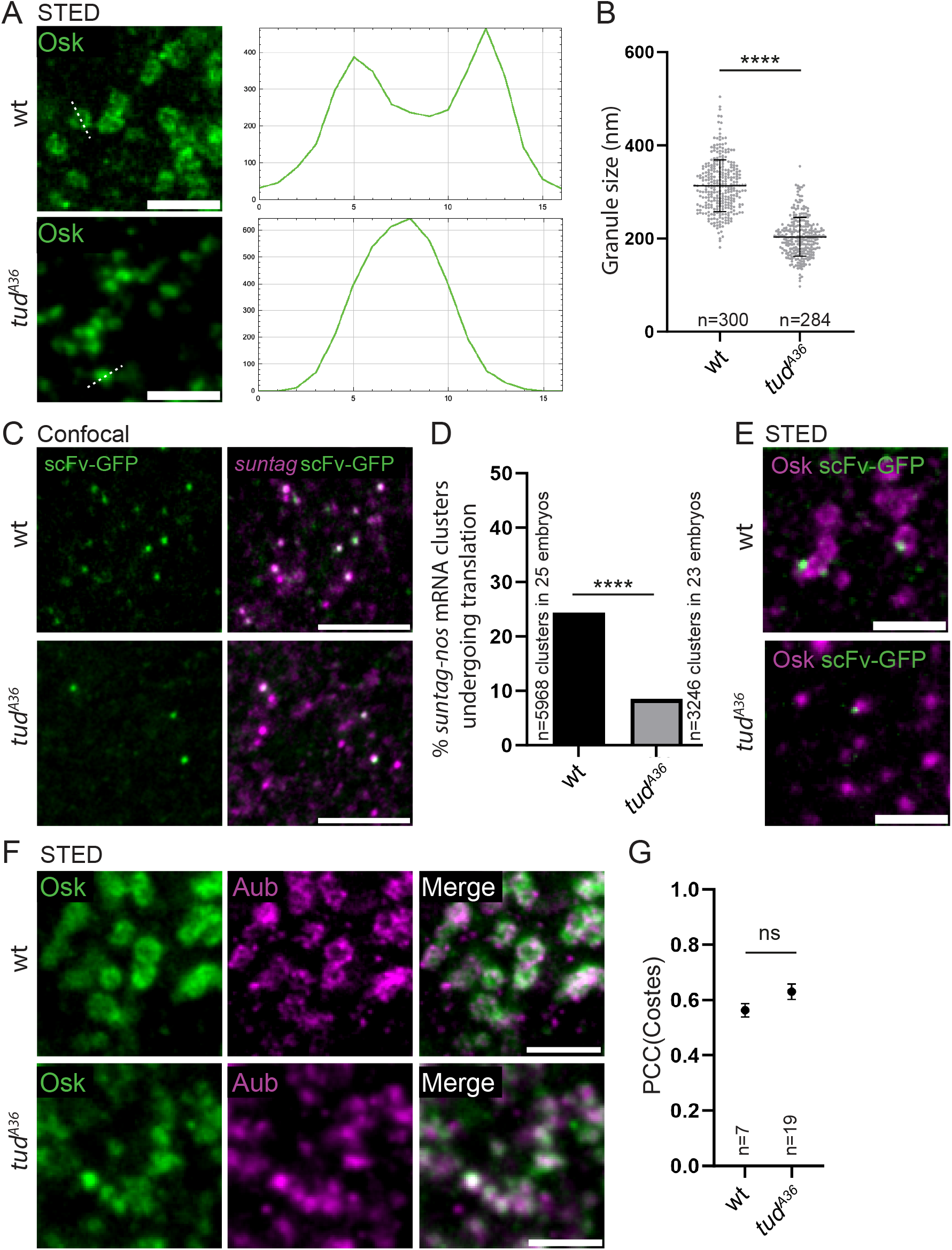
*nos* mRNA translation depends on germ granule biphasic architecture. (**A**) STED images of wild-type and *tud^A36^/Df(2R)Pu^rP133^* (*tud^A36^*) embryos immunostained with anti-Osk antibody. Fluorescence intensity was recorded along the path marked with a white dotted line (right). (**B**) Measurement of germ granule size in wild-type and *tud^A36^/Df(2R)Pu^rP133^* (*tud^A36^*) embryos from images as in (**A**). Horizontal bars represent the mean and SD. **** *p*<0.0001 using unpaired two-tailed Student’s t-test. (**C**) Immuno-smFISH of *suntag-nos/+; scFv-GFP/+* (wild-type) and *suntag-nos tud^A36^/Df(2R)Pu^rP133^; scFv-GFP/+* (*tud^A36^*) embryos with anti-GFP nanobody (green) to reveal scFv-GFP and *suntag* smFISH probe (magenta). (**D**) Percentage of *suntag-nos* mRNA clusters colocalizing with scFv-GFP foci, i.e. undergoing translation, in *suntag-nos/+; scFv-GFP/+* (wild-type) and *suntag-nos tud^A36^/Df(2R)Pu^rP133^; scFv-GFP/+* (*tud^A36^*) embryos from images as in (**C**). **** *p*<0.0001 using the χ^2^ test. (**E**) STED images of immunostaining of *suntag-nos/+; scFv-GFP/+* (wild-type) and *suntag-nos tud^A36^/Df(2R)Pu^rP133^; scFv-GFP/+* (*tud^A36^*) embryos with anti-Osk antibody (magenta) and anti-GFP nanobody (green) to reveal scFv-GFP. (**F**) STED images of immunostaining of wild-type and *tud^A36^/Df(2R)Pu^rP133^* (*tud^A36^*) embryos with anti-Osk (green) and anti-Aub (magenta) antibodies. (**G**) Quantification of colocalization between Osk and Aub using PCC(Costes). Black circles represent the mean and error bars represent SEM. The number of embryos is indicated (n). ns: non-significant using unpaired two-tailed Student’s t-test. Scale bars: 1 µm.

Aub activates *nos* mRNA translation initiation ^43^ and Vasa was also reported to play a role in activating translation ^60^. We, therefore, asked whether Aub or Vasa recruitment to germ granules was impaired in *tud^A36^* embryos, which might contribute to reduced *suntag-nos* mRNA translation. Immunostaining to examine the posterior recruitment of Aub and Vasa together with Osk in *tud^A36^* mutant embryos showed that their recruitment was not affected as quantified using confocal imaging and measuring immunostaining intensity relative to that of Osk (Fig. S10D-G). In addition colocalization of Aub with Osk analyzed using STED microscopy showed that Aub was recruited to germ granules in *tud^A36^* embryos, with a colocalization with Osk similar to that in wild-type embryos (Fig. 6F, G).

We conclude that translation is strongly reduced in *tud^A36^* mutant embryos due to the defective architecture of germ granules, revealing that germ granule core/shell organization plays a critical role in mRNA translation.

## Discussion

In this manuscript, we use *Drosophila* germ granules as a model system to address the importance of the multi-layered organization of biomolecular condensates for their functions. *Drosophila* germ granules ensure two opposite functions: storage of translationally repressed mRNAs and sequential translational activation of the same mRNAs. Using super-resolution STED microscopy, we find that these germ granules have a biphasic organization with an accumulation of protein components in their outer phase. Analyses of mRNA localization within germ granules combined with the suntag approach to record ongoing translation reveal that the storage of translationally repressed mRNAs takes place in the core of germ granules, whereas translation occurs in the shell and at the surface and periphery of the granules. Using 5’end and 3’end probes of the same mRNAs, we show that mRNA orientation within germ granules reflects their translation status, 5’ends of translated mRNAs pointing towards the external regions of germ granules. Moreover, mRNA compaction correlates with their translation levels, translational repression leading to higher mRNA compaction, as it was described recently in stress granules ^61^. Finally, taking advantage of a mutant in which the biphasic architecture of germ granules is lost, we show that their core/shell organization promotes translation. These results demonstrate the functional compartmentalization of germ granules, a process that is central for germ cell specification and development.

We propose a model in which repressed mRNAs are stored in a compacted state in the core of germ granules and translocate to the outer phase for translation. To our surprise, we did not identify translational repressors accumulating in the core of germ granules. Together with the information that untranslated mRNAs have their 5’- and 3’ regions closer than translated mRNAs, and are thus more compacted, these data suggest that translational repression in the core of germ granules would involve a mechanism based on mRNA compaction and compartmentalization away from the translation machinery, independent of specific translational repressors. Nonetheless, several lines of evidence point to a second mechanism involving translational repressors. First, the 3’end of the repressed mRNA, *cycB* is oriented towards the external part of germ granules -outer phase and immediate periphery-, and we also find the association of Smaug translational repressor with the external region of the granules, suggesting that Smaug might bind repressed mRNA 3’UTRs in this location. Smaug interaction with Osk in this external part of germ granules would relieve translational repression ^45,46^. Second, *suntag-nos* translation at germ granules is reduced in *png* mutant embryos and Png kinase is known to relieve translational repression through phosphorylation of translational repressors ^53^. These data suggest the implication of translational repressors in translational regulation at germ granules. Third, consistent with this, translational activation of germ granule mRNAs is sequential, again suggesting the role of specific repressors, or combinations of repressors whose effect would be relieved sequentially to achieve timely translation of specific mRNAs.

A major information from this study is that *Drosophila* germ granules have a function in translation. These granules are not incidental condensates regarding the translation of germ granule mRNAs ^4^ as translation of these mRNAs does not take place anywhere else in the embryo. Therefore, germ granules have an essential function in localized translation of specific mRNAs, which is required for germ cell development. Another key finding is the fact that translation takes place in a defined region of germ granules, the outer phase, indicating that the biophysical properties of this outer phase allow translation. Quantifications of *nos* mRNA levels in germ granules have shown that these levels do not decrease with time up to the formation of primordial germ cells ^62^. This indicates that *nos* mRNAs might be translated several times at germ granules without decay. These consecutive rounds of translation would be possible due to the capacity of the outer phase to promote translation, bypassing the necessity for mRNAs to leave germ granules to be translated, as it is the case in *C. elegans* ^22,23^. Indeed, we found that translation initiation factors concentrate in this phase of germ granules and that the lack of the outer phase in *tud* mutant embryos strongly impairs translation. In addition to a role in localizing translation, germ granules might be involved in increasing translation efficiency through the outer phase properties that would allow consecutive rounds of mRNA translation in the absence of decay.

RNA granules have a recognized role in mRNA storage (e.g. P bodies and stress granules), but their function in translational activation has remained more elusive and is currently an emerging question. A recent study showed that translation can occur in stress granules, although these granules are mostly composed of untranslated mRNAs ^63^. More recently, another study revealed that the formation of RNA granules driven by the FXR1 protein led to the translation of mRNA targets through translation initiation factor recruitment, thus identifying the first RNA condensate specialized in translation activation ^64^. Finally, a recent analysis of zebrafish embryo germ granules reported that mRNA translation required their translocation to the granule periphery ^65^. Here, we found that translation occurs in a specific phase of germ granules, the outer phase, highlighting the functional relevance of RNA granule higher-order organization. Understanding how translation is linked to the formation and biophysical properties of this specific phase of germ granules represents an interesting challenge for future studies.

## Acknowledgments

We are grateful to A. Arkov, M. Lagha, R. Lehmann and the Bloomington *Drosophila* Stock Center for providing *Drosophila* stocks. We thank E. Izaurralde, P. Lasko, A. Nakamura, M. Siomi, A. Vincent and R. Wharton for the gifts of antibodies. We thank C. Jahan for producing the anti-Osk antibody. We are very grateful to E. Bertrand and F. Slimani for their help with image quantification and to the MRI-IGH imaging facility and J.M. Langerak for OMX imaging. This work was supported by the CNRS-University of Montpellier UMR9002, ANR (ANR-19-CE12-0031, ANR-21-CE12-0035-01), MSDAVENIR and FRM (Equipe FRM EQU202303016322). AR held a salary from ANR, Fondation ARC and CNRS, AH held a PhD fellowship from the French Ministry and CG held a salary from ANR and MSDAVENIR.

## Author contributions

Conceptualization: AR, MS

Methodology: AH, AR, MS

Investigation: AH, AR, CG

Visualization: AH, AR, CG

Funding acquisition: MS

Project administration: MS

Supervision: MS

Writing – original draft: AR, MS

Writing – review & editing: AR, MS

## Data and materials availability

All data are available in the main text or the supplementary materials.

## Inclusion and Ethics statement

We support inclusive, diverse and ethical conduct in research.

