## Supplementary material for "Germ granule higher-order organization coordinates their different functions"

**Extended data**

**The PDF file includes:**

Materials and Methods

Supplementary Text

References

Supplementary Figure legends (S1 to S10)

Tables S1 to S5

**Materials and Methods**

***Drosophila lines***

*$w^{1118}$* was used as wild type (wt). Mutant alleles and transgenic lines were: *UASp-GFP-Aub* [1], *nos-Gal4-VP16* [2], *UASp-HA-eIF3d* [3], *Vasa-GFP$^{KI}$*, *GFP-Tud$^{KI}$* and *vasa-tdTom$^{KI}$* [4], *nos-scFv-GFP* [5], *nos$^{BN}$/TM3sb* [6], *bel$^{CC00869}$* [7,8], *png$^{1058}$/FM6a* [9] and *tud$^{A36}$/Cyo* [10]. The *nos-suntag-nos* line was generated in this study by PhiC31 recombination into *attP40* site. The genotypes of embryos (aged 0-2 h or less) indicated throughout were the genotypes of mothers. Females of the indicated genotypes were crossed with wt males.

***Cloning and recombineering***

To produce the *nos-suntag-nos* transgene, the sequence coding for 12 Suntag repeats was amplified by PCR from a clone provided by E. Bertrand [11]. The PCR fragment was cloned after the *nos* start codon into the pBSKS-R5561 plasmid that contains a 5.7 kb *nos* genomic fragment (a gift from R. Wharton [12]), using NEBuilder HiFi DNA Assembly (NEB). The *nos* genomic fragment containing *suntag* sequence was digested by *EcoR*I and *Not*I and cloned into the *pattB* vector (DGRC #1420) digested by *EcoR*I and *Not*I. The resulting plasmid was validated by sequencing and sent for injection (BestGene Inc) to be inserted into the *Drosophila* genome by PhiC31 recombination at the *attP40* site.

***Immunostaining***

0–2 h-embryos were collected in a basket from plates, washed in tap water and dechorionated using commercial bleach for 2 min and rinsed. Embryos were then fixed at the interface of a 1:1 solution of formaldehyde 36% /heptane for 5 min, followed by 100% methanol devitellinization.

Embryos were progressively rehydrated in 75%, 50% and 25% methanol diluted in PBS 0.1% Tween, blocked in 1% BSA for 1 h for confocal imaging or 10% BSA for 3 h for STED imaging, and incubated overnight with primary antibodies. Secondary antibody incubation, after washes in PBS 0.1% Tween, was performed for 1 h at room temperature. Embryos were mounted in Vectashield (Vector Laboratories) for confocal imaging or Abberior liquid anti-fade (Abberior) for super-resolution imaging (STED, OMX and Airyscan). For super-resolution imaging the posterior pole of embryos was sliced using a thin needle in mounting medium and mounted posterior side up (Fig. S1A). For stage 14 oocytes, ovaries were dissected from well fed females in Schneider medium and fixed in hypotonic solution (1:1 heptane/PBS 1X, 50 mM EGTA pH8, 10.24% formaldehyde) to prevent egg activation, for 1.5 h at room temperature. Then ovaries were rinsed three times in PBS 0.1% Tween, transferred to a dissection dish where oocytes were dissociated from the rest of the ovary by multiple pipetting. Oocytes were put on a frosted slide and rolled with another frosted slide to remove chorion and vitelline membrane. Oocytes were transferred in a tube and processed for immunostaining as described for early embryos. Primary antibodies used were: mouse anti-Aub (1/1000, clone 4D10, a gift from M. Siomi [13]), rabbit anti-CCR4 (1/200) [14], rabbit anti-Cup (1/1000, a gift from R. Wharton, [15]), rabbit anti-Me31B (1/2000, a gift from A. Nakamura, [16]), rabbit anti-Nos (1/1000, a gift from A. Nakamura), mouse anti-Not1 (1/100, clone 2G5, [14]), rabbit anti-Osk (1/1000, this study), rabbit anti-PABP (1/500, a gift from A. Vincent), rabbit anti-Smg (1/1000, [17]), rabbit anti-Tud (1/500, a gift from P. Lasko, [18]), mouse anti-HA (1/2000, ascites produced from clone 12CA5), rat anti-Vasa (1/50, Developmental Studies Hybridoma Bank), rabbit anti-GFP (1/1000, Invitrogen), and mouse anti-GFP (1/1000, Roche). Secondary antibodies used were: FluoTag®-X4 anti-GFP nanobodies StarRed or Atto488 (1/500, NanoTag Biotechnologies), goat anti-rabbit IgG Alexa-488 (1/800, Invitrogen), donkey anti-rabbit

IgG Cy5 (1/1000, Jackson ImmunoResearch), goat anti-mouse IgG StarRed (1/1000, Abberior), goat anti-rabbit IgG StarRed (1/1000, Abberior), and goat anti-rabbit IgG Star580 (1/1000, Abberior).

***smFISH and immuno-smFISH***

smFISH on 0-2 h-embryos were performed as previously described [3] with smFISH probes from Stellaris. For immuno-smFISH, rehydrated embryos were post-fixed in 4% formaldehyde for 20 min, rinsed three times for 10 min in PBS 0.1% Tween and processed for smFISH followed by immunostaining. smFISH on stage 14 oocytes were performed as previously described [19] using 10% formamide. Probe sequences for *nos*, *gcl*, *pgc* and *cycB* are listed in [20]. Probe sequences to detect *suntag*, *nos* 5'end, *nos* 3'end, *cycB* 5'end and *cycB* 3'end are listed in Supplementary Tables S1 to S5.

***Puromycin treatment***

To treat embryos with puromycin, we adapted the permeabilization protocol previously described [21]. Dechorionized embryos were put in a 1:1 solution of DL-limonene (Sigma)/heptane with a 100 µl drop of PBS 1X containing 2 mg/ml puromycin (Sigma). Puromycin was initially diluted in water at 50 mg/ml. For control embryos, the puromycin solution was replaced by water alone. Embryos were shaken at maximum speed on a rotating plate for 40 min, rinsed in heptane and processed for immunostaining.

***Generation of anti-Osk antibody***

The open reading frame of *osk* (short isoform) without the start codon was cloned after the *GST* sequence into the *pGEX-4T-1* vector to express a GST-Osk fusion protein in BL21 *E. coli* bacteria. The GST-Osk fusion protein was isolated on an acrylamide gel, purified using columns (Amicon) and injected into rabbits by Agro-Bio. The polyclonal rabbit antibody against this fusion protein was validated using immunostaining in ovaries and embryos.

***Microscopy***

Confocal microscopy was performed using a Leica SP8 confocal scanning microscope with objectives 20X Plan Apochromat 0.75 NA Imm Corr for whole embryos (Fig. S3C) or 63X Plan Apochromat 1.4 NA oil DIC for other confocal images. STED microscopy was performed using an Abberior STED super-resolution microscope controlled by Imspector software (Abberior Instruments) using a 100X Plan SuperApochromat 1.4 Oil objective. Excitation and depletion laser powers were adjusted according to the strength of the signal to optimize the resolution without bleaching the sample. OMX microscopy was performed by Deltavision OMX run by SoftWoRx software using a 100X Plan SuperApochromat 1.4 Oil (DIC) objective. Image quality was verified using SIMcheck [22]. Airyscan microscopy was performed using confocal Zeiss LSM980 AiryScan II 8Y run by Zeiss Zen Blue software using a 63X Plan Apo oil 1.4NA objective.

***Image analyses and quantifications***

*Quantification of mRNA molecule numbers.* mRNA content in germ granules was quantified using FISH-quant [11,23]. Briefly, the mean intensity of single mRNA molecule was calculated from the signal of single particles in the soma as they mostly correspond to single mRNA molecules (83% of single particles for *nos* mRNA [24]). To determine the number of mRNA molecules present in each mRNA cluster at the posterior pole, the integrated intensity of each cluster was divided by the integrated intensity of the mean single mRNA molecules.

*Quantification of germ granule size and shell size.* On STED acquisitions of Osk immunostaining, a line was drawn through the diameter of each granule and intensities along the line were measured using FIJI plot profile, giving two peaks that reflected the donut-like shape of germ granules (Fig. 1A). To determine the limit of the signal, the peak height was divided by half and used to define the signal edges, allowing the measurement of the granule diameters and shell thickness.

*Quantification of the localization of mRNAs or scFv-GFP foci within the germ granule biphasic structure.* To analyze the localization of mRNA foci (visualized either with the full-length, the 5'end, or the 3'end probes) or scFv-GFP foci within germ granule biphasic organization, we first applied a Laplacian Gaussian filter on the channel showing the germ granule staining to define the signal edges and thus the compartments (core, shell, periphery). Then mRNA coordinates were determined using RS-FISH ImageJ pluggin [25]. A line was drawn across the granule and mRNA foci and pixel intensities along the line were recorded on Fiji Plot profile. The distance between the peak of the shell and the peak of the mRNA focus was measured on the plot. If the mRNA focus was located toward the external part of the granule, the measure was given a positive value. If the mRNA focus was toward the internal part of the granule, it was given a negative value. As the size of the shell is not identical between granules, the shell size was also measured for each measurement of the distance between the shell peak and mRNA focus, allowing us to determine the position of the mRNA focus within the granule using the following ratio:

distance between the peak of mRNA focus and the peak of the shell / (shell size/2).

If the ratio was >1, the RNA focus was classified as “periphery”. If the ratio was ≤ -1, the RNA focus was classified as “core”. If the ratio was >-1 and ≤1, the RNA focus was classified as “shell”.

Quantifications of these ratios were shown as graphs, and the localization of mRNA foci were represented on hypothetic granules using radar plots. The same method was applied to quantify the position of scFv-GFP foci within germ granules.

*Analysis and quantification of colocalization of scFv-GFP foci with suntag mRNA clusters and/or germ granules.* On confocal images, germ granules appear as full dots or foci. To measure the association of scFv-GFP foci with germ granules or *suntag* mRNA clusters, we used FIJI plugin ComDet v.0.5.5 (https://github.com/UU-cellbiology/ComDet) to define the different foci and measure their colocalization. To visualize the colocalization of scFv-GFP foci with *suntag* mRNA clusters at germ granules (i.e. translation at germ granules, Fig. S3C), we segmented the signals from *suntag* smFISH and scFv-GFP and created a mask that showed their colocalization (translation foci) using FIJI Image Calculator function. We then overlaid this mask on the channel showing germ granules visualized with anti-Osk immunostaining and quantified the colocalization of translation foci with germ granules.

*Distance and colocalization between mRNA 5'end and 3'end.* To measure the distance between *nos* mRNA 5'end and 3'end probes, the 3'end signal was segmented and defined as ROI. 5'end foci were identified using FIJI plugin ComDet v.0.5.5 (https://github.com/UU-cellbiology/ComDet) and their center was mapped by multipoint selection and added as ROI. The minimal distance between the edge of the 3'end ROI and the center of the 5'end foci was calculated using a FIJI plugin developed previously [26]. Colocalization corresponds to a distance of 0.

*Distance between Smaug foci and germ granules.* To analyze the distance between Smaug foci and germ granules, Smaug signals above threshold were identified using the Fiji Find Maxima function. The signal for the granule corresponding to anti-Osk immunostaining was segmented and defined as ROI. The minimal distance between the edge of the granule ROI and the center of

Smaug foci was calculated using a FIJI plugin developed previously [26]. As a control (Smg tilted), Smaug channel was horizontally and vertically rotated and the same measurement was performed.

*Protein colocalization within germ granules*. For images acquired with STED imaging, the degree of colocalization between signals corresponding to two immunostaining was quantified using PCC(Costes). The PCC method determines a threshold based on the mean fluorescence intensity for each signal. Then, the method analyzes each pixel and evaluates if signals are above or below their respective threshold, increasing PCC value if both signals go to the same direction, decreasing its value if they go to opposite directions. PCC ranges from 1 that indicates perfect colocalization, to -1 that indicates exclusion. To validate the significance of this colocalization, the PCC(Costes) value was calculated for each image. Briefly, images were randomized by shuffling pixels, the PCC value was calculated and compared to the original image. This process was repeated 200 times to evaluate the significance of the original picture. Costes *p*-value ranges from 0 to 1, where 0 indicates random colocalization and 1 indicates significant colocalization.

*Quantification of germ granule main component levels.* To quantify the level of Aub and Vasa in $tud^{A36}$ mutants, ROIs delimiting the germ plasm on maximum Z projection of 40 planes from confocal images were defined. The fluorescence integrated density of Aub or Vasa was measured and normalized to Osk fluorescence integrated density levels.

#### *Statistical analyses*

Statistical tests were performed using GraphPad. For each figure, the tests are indicated in the figure legend.

### Supplementary Text

#### *Imaging germ granule core/shell organization*

*Drosophila* embryos have an ovoid shape and the germ plasm is only accessible by imaging deep into the sample, which decreases resolution. Therefore, to reach the resolution required to resolve germ granule organization, the distance between the sample and the objective had to be reduced. This was achieved using two approaches: 1) either optimizing the mounting of embryos such that the posterior pole was directly facing the coverslip, or 2) slicing the posterior of embryos. In the first approach, embryos were mounted in a high concentration, making embryos pile up one upon another. Only embryos with the posterior pole facing up were imaged. In the second approach, the posterior of embryos was sliced using a needle and mounted with the posterior side up (Fig. S1A). Both approaches led to a level of resolution allowing to reveal germ granule structure. The requirement of these mounting methods was illustrated by imaging germ granule using the Airyscan system. This is a confocal laser scanning set up where an array of detectors allows the improvement of resolution up to 120 nm, giving super-resolution-like images. Using whole mount embryos, we could not observe germ granule core/shell organization (Fig. S1C, top). However, using sliced posterior poles, we easily observed GFP-Aub donut-like shapes, even with GFP fluorescence, in the absence of antibody staining (Fig. S1C, bottom). The germ granule core/shell architecture was observed using three different microscopy techniques, STED, Airyscan and OMX, with and without antibody staining, establishing this biphasic organization.

### Supplementary Figure legends

**Fig. S1. Biphasic organization of *Drosophila* germ granules visualized with three super-resolution microscopy techniques.**

(**A**) Illustration of the mounting of the embryo posterior following slicing. (**B**) Immunostaining of *UASp-GFP-Aub; nos-Gal4* embryos with anti-GFP antibody, showing the same germ granules imaged using confocal (top) or STED super-resolution (bottom) microscopy. Fluorescence intensity (right) was recorded along the path marked with a red line. (**C**) Airyscan imaging of *UASp-GFP-Aub; nos-Gal4* embryos in whole mount (top) or as a sliced posterior pole (bottom). Fluorescence intensity (right) was recorded along the path marked with a red line. GFP fluorescence was directly recorded without antibody staining. (**D**) Quantification of colocalization between the indicated components shown in Fig. 1A, using PCC(Costes). Black circles represent the mean and error bars represent SEM. The number of embryos is indicated (n). (**E**) 3D-OMX imaging of an *UASp-GFP-Aub; nos-Gal4* sliced posterior pole. YZ and XZ show the orthogonal views of the acquisition. GFP fluorescence was directly recorded without antibody staining. Scale bars: 1 µm.

**Fig. S2. Defective germ granule organization in the absence of Aub.**

(**A**) Immunostaining of *UASp-osk-bcd3'UTR/nos-Gal4* (*osk-3'bcd*) embryos with anti-Osk antibody showing Osk protein localization at both poles using confocal microscopy (top). *osk-bcd3'UTR* chimeric mRNA is recruited to the embryo anterior pole due to the presence of *bicoid* (*bcd*) 3'UTR. Visualization of germ granule organization using STED microscopy (bottom) showing that both anterior and posterior germ granules are biphasic. Fluorescence intensity was recorded along the path marked with a red dotted line. (**B**) Immunostaining of *aub*$^{QC42/HN2}$; *UASp-*

*osk-bcd3'UTR/nos-Gal4* (*aub$^{-/-}$; osk-3'bcd*) embryos with anti-Osk antibody showing Osk protein localization only at the anterior pole using confocal microscopy (top). Visualization of germ granule organization using STED microscopy (bottom) showing the loss of germ granule biphasic structure. Fluorescence intensity was recorded along the path marked with a red dotted line. (**C**) Quantification of germ granule size in *UASp-osk-bcd3'UTR/nos-Gal4* (*osk-3'bcd* Posterior and Anterior) and *aub$^{QC42/HN2}$*; *UASp-osk-bcd3'UTR/nos-Gal4* (Anterior *aub$^{-/-}$*) embryos. Horizontal bars represent the mean and SD. ns: non-significant, **** $p$<0.0001 using the unpaired two-tailed Student's t-test. The number of granules is indicated (n). (**D**) STED imaging of immuno-smFISH of *UASp-osk-bcd3'UTR/nos-Gal4* (*osk-3'bcd*) and *aub$^{QC42/HN2}$*; *UASp-osk-bcd3'UTR/nos-Gal4* (*aub$^{-/-}$; osk-3'bcd*) embryos with anti-Osk antibody and *nos* smFISH probe showing the lack of *nos* mRNA in germ granules from *aub$^{QC42/HN2}$* mutant embryos. Scale bars: 50 µm in (**A**, **B**, top), 1 µm in (**A**, **B**, bottom and **D**).

**Fig. S3. Visualization of *nos* translation at germ granules using *suntag-nos*.**

(**A**) Schematic representation of the *suntag-nos* construct. (**B**) Z-projection of confocal images of smFISH on wild-type (top) and *suntag-nos/+* (bottom) embryos hybridized with *nos* and *suntag* probes, respectively. The corrected total fluorescence intensity at the posterior pole and in the whole embryo was measured to assess the percentage of mRNA localized at the posterior pole. (**C**) Immuno-smFISH of *suntag-nos/+; scFv-GFP/+* embryos with anti-Osk antibody (grey), anti-GFP (green) nanobody to reveal scFv-GFP, and smFISH *suntag* probes (magenta). Colocalization of *suntag* and scFv-GFP signals (cyan) indicates ongoing *suntag-nos* translation. (**D**) Quantification using FISH-quant of the number of *nos* and *suntag-nos* mRNA molecules per cluster at the posterior pole from confocal images of smFISH on wild-type and *suntag-nos* embryos, respectively. Horizontal bars represent the mean and SD. **** $p<0.0001$ using the unpaired two-tailed Student's t-test. The number of mRNA clusters is indicated (n). (**E**, **F**) Three-color STED imaging of immuno-smFISH of *suntag-nos/+; scFv-GFP/+* embryos with anti-Osk antibody (grey), anti-GFP nanobody (green) to reveal scFv-GFP and smFISH *suntag* probes (magenta). Scale bars: 50 µm in (**B**), 1 µm in (**C**, **E**, **F**).

**Fig. S4. Localization of translation initiation factors and translational repressors in the germ plasm.**

(**A, B**) Localization of translation intiation factors. STED imaging of immunostaining of *UASp-GFP-Aub/UASp-HA-eIF3d; nos-Gal4/+* (**A**) and *UASp-GFP-Aub; nos-Gal4* (**B**) embryos with anti-GFP (magenta) to detect Aub, anti-HA (green) to detect eIF3d (**A**) and anti-PABP (green) (**B**). (**C-I**) Localization of translational repressors. STED imaging of immunostaining of *UASp-GFP-Aub; nos-Gal4*, *GFP-belle* and wild-type embryos with antibodies to reveal the indicated proteins. Germ granule markers are in magenta and translational repressors are in green. A larger germ granule exemplifying those present at later stages in primordial germ cells (PGC) is shown in (**F**). (**J**) Quantification of colocalization of GFP-Aub or Osk with the indicated proteins using PCC(Costes) from images as in (**A-E** and **G-I**). Black circles represent the mean and error bars represent SEM. The number of embryos is indicated (n). (**K**) Quantification of the distance between Smaug (Smg) foci and the edge of germ granules marked with GFP-Aub from images as in (**C**). Vertical bars represent the mean and SD. ** $p<0.01$ using the $\chi^2$ test. Scale bars: 1µm.

**Fig. S5. Schematic representation of smFISH probes.**

The boxes represent oligos composing the probes. Grey boxes indicate probes covering full length mRNAs or coding sequences; orange boxes indicate probes covering mRNA 5’ends; and green boxes indicate probes covering mRNA 3’ends.

**Fig. S6. Decrease of translation in *png* mutant embryos.**

(**A**) Fluorescent confocal imaging of smFISH on *suntag-nos/+; scFv-GFP/+* (wt) and $png^{1058/1058}$; *suntag-nos /+; scFv-GFP/+* ($png^{-/-}$) embryos, showing scFv-GFP (green) and *suntag-nos* mRNA (magenta). (**B**) Percentage of *suntag-nos* mRNA clusters undergoing translation in *suntag-nos/+; scFv-GFP/+* (wt) and $png^{1058/1058}$; *suntag-nos /+; scFv-GFP/+* ($png^{-/-}$) embryos. The graph represents the percentage of *suntag-nos* mRNA clusters colocalizing with scFv-GFP foci from images as in (**A**). **** $p<0.0001$ using the $\chi^2$ test. (**C**) Quantification using FISH-quant of the number of *suntag-nos* mRNA molecules per cluster at the posterior pole from confocal images of smFISH on *suntag-nos/+; scFv-GFP/+* (wt) and $png^{1058/1058}$; *suntag-nos /+; scFv-GFP/+* ($png^{-/-}$) embryos from images as in (**A**). Horizontal bars represent the mean and SD. **** $p<0.0001$ using the unpaired two-tailed Student’s t-test. The number of mRNA clusters is indicated (n). Scale bars: 5 μm.

**Fig. S7. *cycB* mRNA localization and compaction are not affected in *png* mutant embryos.**

(**A**) STED imaging of immuno-smFISH of wild-type and $png^{1058}$ mutant ($png^{-/-}$) embryos with anti-Osk antibody and *cycB* smFISH probe. (**B**) Percentage of localization of *cycB* mRNA foci in wild-type and $png^{1058}$ mutant embryos, in the core (black), in and at the surface of the shell (grey) and at the immediate periphery (white) of germ granules from images as in (**A**). ns: non-significant using the $\chi^2$ test. (**C**) STED images of smFISH against *cycB* 5’end (5'-*cycB*, magenta) and 3’end

(3'-*cycB*, green) in wild-type and $png^{1058}$ mutant embryos. (**D**) Measurement of the distance between *cycB* 3'end foci and the edge of *cycB* 5'end foci in wild-type and $png^{1058}$ embryos from images as in (**C**). The histogram shows the percentage of *cycB* 3'end foci in each distance class. (**E**) Percentage of *cycB* 3'end foci colocalizing with *cycB* 5'end foci in wild-type and $png^{1058}$ embryos. ns: non-significant using the $\chi^2$ test. Scale bars: 1 µm.

**Fig. S8. Illustration of the method to measure the distance between 5'end and 3'end smFISH probes.**

The most internal signal, here *3'-nos* in green, was segmented to define regions of interest (ROI) that corresponded to the area of the probe signal. The signal from *5'-nos* smFISH probes in magenta was segmented as foci. The distance between the edge of the ROI defined by *3'-nos* and the center of *5'-nos* foci was then measured. When *5'-nos* foci were within *3'-nos* ROI, the measured distance was 0.

**Fig. S9. *suntag-nos* is not translated in stage 14 oocytes.**

Confocal image of $vasa\text{-}tdTom^{KI}$*/suntag-nos; scFv-GFP/+* stage 14 oocyte germ plasm. GFP and tdTomato were recorded without antibody staining. The GFP fluorescence was absent in 100% of stage 14 oocytes (n=14). Scale bar: 5 µm.

**Fig. S10. Characterization of germ granule content in $tud^{A36}$ mutant.**

(**A**) Schematic representation of Tud protein and the point mutation in $tud^{A36}$ mutant [10]. Boxes represent the eleven Tudor domains. The mutation in $tud^{A36}$ is in the first Tudor domain. (**B**) Quantification using FISH-quant of the number of *suntag-nos* mRNA molecules per cluster at the posterior pole from confocal images of smFISH on *suntag-nos/+; scFv-GFP/+* (wt) and *suntag-nos* $tud^{A36}$*/Df(2R)Pu*$^{rP133}$*; scFv-GFP/+* ($tud^{A36}$) embryos from images as in Fig. 6C. Horizontal bars represent the mean and SD. **** $p$<0.0001 using the unpaired two-tailed Student's t-test. The number of mRNA clusters is indicated (n). (**C**) Percentage of scFv-GFP foci in *suntag-nos/+; scFv-GFP/+* (wild-type) and *suntag-nos* $tud^{A36}$*/Df(2R)Pu*$^{rP133}$*; scFv-GFP/+* ($tud^{A36}$) embryos, localized in the core (black), the shell (grey) and at the immediate periphery (white) of wild-type germ granules, and in the monophasic germ granule (pink) and at their immediate periphery (white) in $tud^{A36}$ embryos. The striped pink rectangle represents the percentage of scFv-GFP foci within 50 nm of the granule surface. (**D**) Confocal images of immunostaining of wild-type (wt) and $tud^{A36}$*/Df(2R)Pu*$^{rP133}$ ($tud^{A36}$) embryos with anti-Aub (magenta) and anti-Osk (green) antibodies. (**E**) Quantification of Aub protein levels (signal intensity) normalized to Osk levels. ns: non- significant using the unpaired two-tailed Student's t-test. (**F**) Confocal images of immunostaining of wild-type (wt) and $tud^{A36}$*/Df(2R)Pu*$^{rP133}$ ($tud^{A36}$) embryos with anti-Vasa (magenta) and anti-Osk (green) antibodies. (**G**) Quantification of Vasa protein levels (signal intensity) normalized to Osk levels. ns: non-significant using the unpaired two-tailed Student's t-test. Scale bars: 10 µm.

### **Table S1. List of smFISH probes to detect the *suntag* sequence.**

| Probe sequence | Probe Name |
|---|---|
| ggtggtagttcttgctcag | suntag_1 |
| ctacccttcttcagtctgg | suntag_2 |
| gctcaaaagttcttcaccg | suntag_3 |
| gccacttcgttctcaagat | suntag_4 |
| gccagaacctttcttaaga | suntag_5 |
| tgaaagcagttcttctcca | suntag_6 |
| tcattttccaggtggtaat | suntag_7 |
| cccttttcagtctagcta | suntag_8 |
| atttttgctcagcaactcc | suntag_9 |
| tgctacttcgttctccaaa | suntag_10 |
| atccggaccctttctttag | suntag_11 |
| tcgagagtaactcctcacc | suntag_12 |
| gccacttcgttttcgagat | suntag_13 |
| accactgcccttttttagc | suntag_14 |
| agataatagctcttctcca | suntag_15 |
| attttcgaggtggtagttt | suntag_16 |
| cttttttaagcgtgccacc | suntag_17 |
| accactgccactagtactt | suntag_18 |
| attcttggatagtagctct | suntag_19 |
| gctacctcgttctcaagat | suntag_20 |
| ccggaacccttcttcaaac | suntag_21 |
| agttcttcgagagcagttc | suntag_22 |
| gcgacctcattttcaagat | suntag_23 |
| cccgatcccttttttaatc | suntag_24 |
| gaaagtagttcctcaccac | suntag_25 |
| cttcgttttcgaggtggta | suntag_26 |
| ccctgaacctttctttaat | suntag_27 |
| actcagtaattcttcaccc | suntag_28 |
| gctacctcattttccagat | suntag_29 |
| aaccactccccttttttag | suntag_30 |
| ttcgatagcaattcctcgc | suntag_31 |
| agcaacttcgttctcaaga | suntag_32 |
| taccagagccctttttgag | suntag_33 |
| tacttagtaattcctcacc | suntag_34 |
| cttgctacttcattctcta | suntag_35 |
| gagcctgagccctttttta | suntag_36 |

**Table S2. List of smFISH probes to detect *nos* 5’end (*5′-nos*).**

| Probe sequence | Probe Name |
|---|---|
| aagctacgcgccaactaa | 5'-nos_1 |
| ttccaggaattttgtggt | 5'-nos_2 |
| aactgcgaagcgtacggc | 5'-nos_3 |
| tcgtatgtcccttagaca | 5'-nos_4 |
| aatcgtgacgcagaggca | 5'-nos_5 |
| ctaaactcgcttttgggt | 5'-nos_6 |
| ccacaaatcctcacccaa | 5'-nos_7 |
| ggttatcgcgcactctac | 5'-nos_8 |
| ggcgaaaatccgggtcga | 5'-nos_9 |
| ccaagttgctgcggaaca | 5'-nos_10 |
| aagttatctgctgctgcg | 5'-nos_11 |
| cctcctctggcgtgaaaa | 5'-nos_12 |
| tgcaggcccagaatgttg | 5'-nos_13 |
| cccactggtatccaaata | 5'-nos_14 |
| aagtggccgacgagttgg | 5'-nos_15 |
| ggcgtaatgggcggactc | 5'-nos_16 |
| agacgtcgacgggtcagg | 5'-nos_17 |
| ggaagtgcgtcgactgcg | 5'-nos_18 |
| gcgctgtcggccagaaaa | 5'-nos_19 |
| ggagcgaattggcggtgg | 5'-nos_20 |
| tggtactgtcgctgcata | 5'-nos_21 |
| tgctggagcagcaagtgg | 5'-nos_22 |
| catggccagttgctgctg | 5'-nos_23 |
| cagcgccaattggtgctg | 5'-nos_24 |
| tttgctggtgactcgcac | 5'-nos_25 |

**Table S3. List of smFISH probes to detect *nos* 3'end (*3'-nos*).**

| Probe sequence | Probe Name |
|---|---|
| tctatctatctggttaacccag | 3'-nos_1 |
| caggcgctatttaaacgttact | 3'-nos_2 |
| aatctctttaaaatcgaacgcg | 3'-nos_3 |
| aagatctataggcacgggataa | 3'-nos_4 |
| tgatcgttcgttgtctatacta | 3'-nos_5 |
| ttcttgaattattgacttggat | 3'-nos_6 |
| caaaattagtttccctttcaca | 3'-nos_7 |
| acgatattgtaagtcttcttta | 3'-nos_8 |
| gccacgacgattgaacaagtat | 3'-nos_9 |
| ttcggattgtaagatatttcta | 3'-nos_10 |
| cagaccaattccattcatcaac | 3'-nos_11 |
| tttacgaaatgaaggcgaccag | 3'-nos_12 |
| atatatcgaaatttttcggccg | 3'-nos_13 |
| attcaaagtgttcctttttcaa | 3'-nos_14 |
| aatgatacgattgacagttcga | 3'-nos_15 |
| tcctttagcaagatttaaattt | 3'-nos_16 |
| cgacgaaagtgttccttgctat | 3'-nos_17 |
| attttacaatgaatgcgtagcc | 3'-nos_18 |
| agtgcggaatgtcaaaatttaa | 3'-nos_19 |
| atactcttcgcttatctatcaa | 3'-nos_20 |
| gtgttgaaatgaatacttgcga | 3'-nos_21 |
| aattatataatgctggcggttg | 3'-nos_22 |
| tcagaatatgtgtacacatttt | 3'-nos_23 |
| tcgagccattgaatttttcatt | 3'-nos_24 |
| tgtaaccatttctttatttggc | 3'-nos_25 |

**Table S4. List of smFISH probes to detect *cycB* 5'end (*5′-cycB*).**

| Probe sequence | Probe name |
|---|---|
| gctgccgtttgaatttga | 5'-cycB_1 |
| ttgcacacgaagcgaggc | 5'-cycB_2 |
| ctccgaaaacctgatcga | 5'-cycB_3 |
| tcgagtgcgggattgtca | 5'-cycB_4 |
| cgcttgtttagctgttga | 5'-cycB_5 |
| gatcgcgtttctgtgacc | 5'-cycB_6 |
| ggctatcacttggtttgg | 5'-cycB_7 |
| cgaagataggcagacgct | 5'-cycB_8 |
| tgcttcttttctatctgt | 5'-cycB_9 |
| ttgtgcccaccattttga | 5'-cycB_10 |
| atcgccacgcattttcag | 5'-cycB_11 |
| agttctccgaagcgttct | 5'-cycB_12 |
| tcttcaattgcacttgct | 5'-cycB_13 |
| ccatggaaggaaccgtca | 5'-cycB_14 |
| gcgcgttttgttgttgcc | 5'-cycB_15 |
| ctgcaaatcgcccaaggc | 5'-cycB_16 |
| gacgacttatgccgcgat | 5'-cycB_17 |
| gcatccttcgctgcgatg | 5'-cycB_18 |
| ccttggagtctttctgtg | 5'-cycB_19 |
| gcgtctgtgagcttgaga | 5'-cycB_20 |
| agctttggcattgcgcag | 5'-cycB_21 |
| agtggctgtttcttccag | 5'-cycB_22 |
| cattgccattgccattgg | 5'-cycB_23 |
| ttgaccttgggcggaacg | 5'-cycB_24 |
| aaaaacgccgacacgccc | 5'-cycB_25 |

**Table S5. List of smFISH probes to detect *cycB* 3’end (*3′-cycB*).**

| Probe sequence | Probe name |
|---|---|
| ttcagcttggcctgaggag | 3'-cycB_1 |
| ggtacttgttgtagatggc | 3'-cycB_2 |
| gcgatcttctggaacttgc | 3'-cycB_3 |
| acaatcgagtccatcagcg | 3'-cycB_4 |
| tatttcctctggctctggc | 3'-cycB_5 |
| aaactaggtaagtcagggtct | 3'-cycB_6 |
| cgcatttttaacgaacacga | 3'-cycB_7 |
| gcaatgcgactacggtaacta | 3'-cycB_8 |
| atgggtgatgcaggtaaagat | 3'-cycB_9 |
| ccagcagattacgatatctta | 3'-cycB_10 |
| agtagccgaaaactgctcaag | 3'-cycB_11 |
| tagtcttaccggaggaactca | 3'-cycB_12 |
| gagatggtgacccagcgaaag | 3'-cycB_13 |
| aaatgaactcgttgcccactc | 3'-cycB_14 |
| aattcttggagagacaacgcc | 3'-cycB_15 |
| tgaaatgatttcacgggtaca | 3'-cycB_16 |
| attagtgataggtaggtaggg | 3'-cycB_17 |
| tattcattcccatcgaaacta | 3'-cycB_18 |
| attttgcacaattttttggat | 3'-cycB_19 |
| ttttatgcgatttatgggaat | 3'-cycB_20 |
| ttacaaatagtctacgtctct | 3'-cycB_21 |
| tgtttttgtatgaatgtgcga | 3'-cycB_22 |
| ttttaatcgttgcttagtggt | 3'-cycB_23 |
| tttgggaaacatcagttagtt | 3'-cycB_24 |
| tctatgtattgttcagagaca | 3'-cycB_25 |

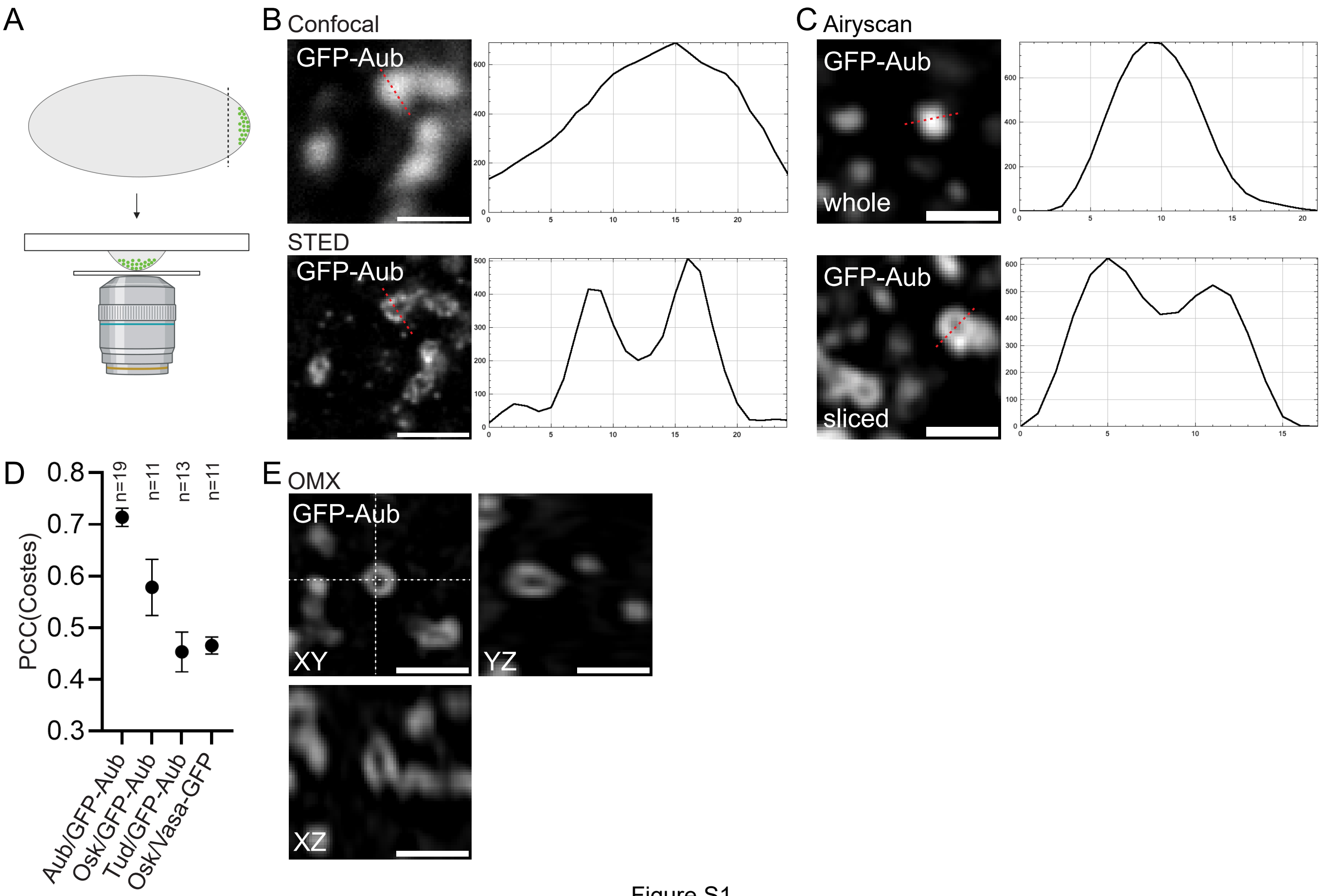
A
B Confocal
GFP-Aub
STED
GFP-Aub
C Airyscan
GFP-Aub
whole
GFP-Aub
sliced
D
PCC(Costes)
n=19
n=11
n=13
n=11
Aub/GFP-Aub
Osk/GFP-Aub
Tud/GFP-Aub
Osk/Vasa-GFP
E OMX
GFP-Aub
XY
YZ
XZ

Figure S1

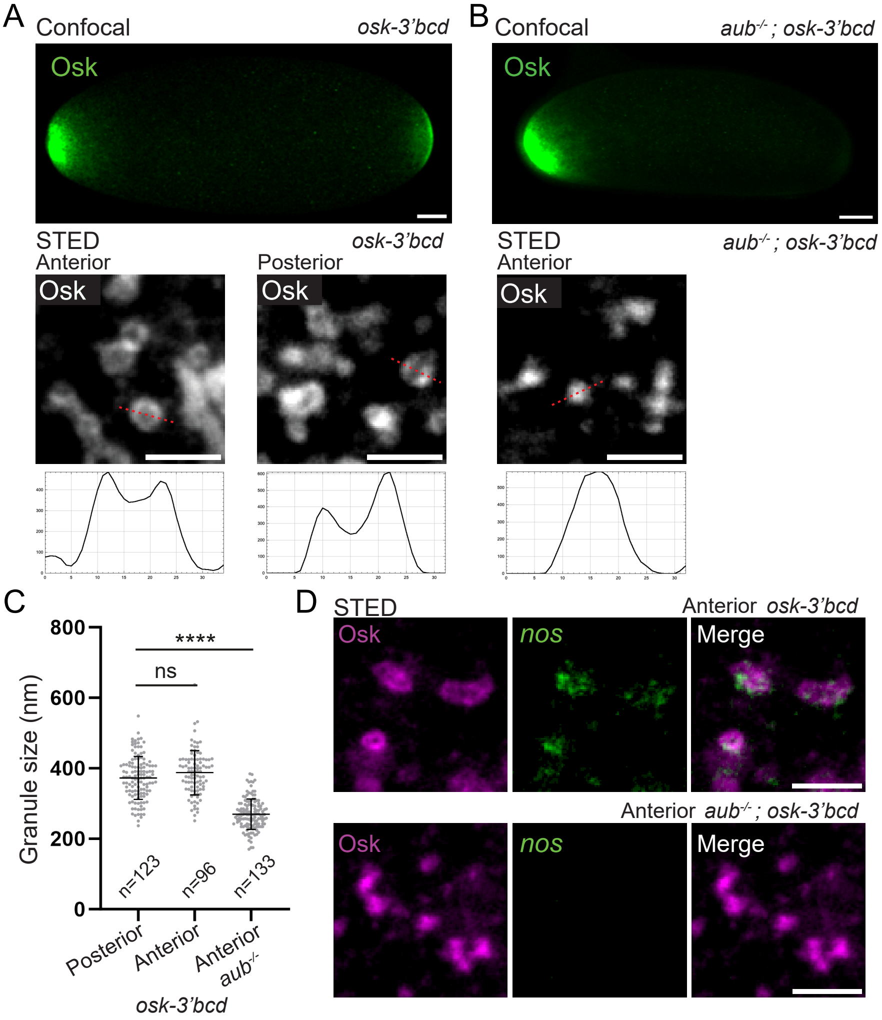
A
Confocal
osk-3'bcd
Osk
STED
Anterior
Posterior
B
aub-/- ; osk-3'bcd
C
Granule size (nm)
800
600
400
200
0
****
ns
n=123
n=96
n=133
Posterior
Anterior
Anterior aub-/-
D
Anterior osk-3'bcd
nos
Merge
Anterior aub-/- ; osk-3'bcd

Figure S2

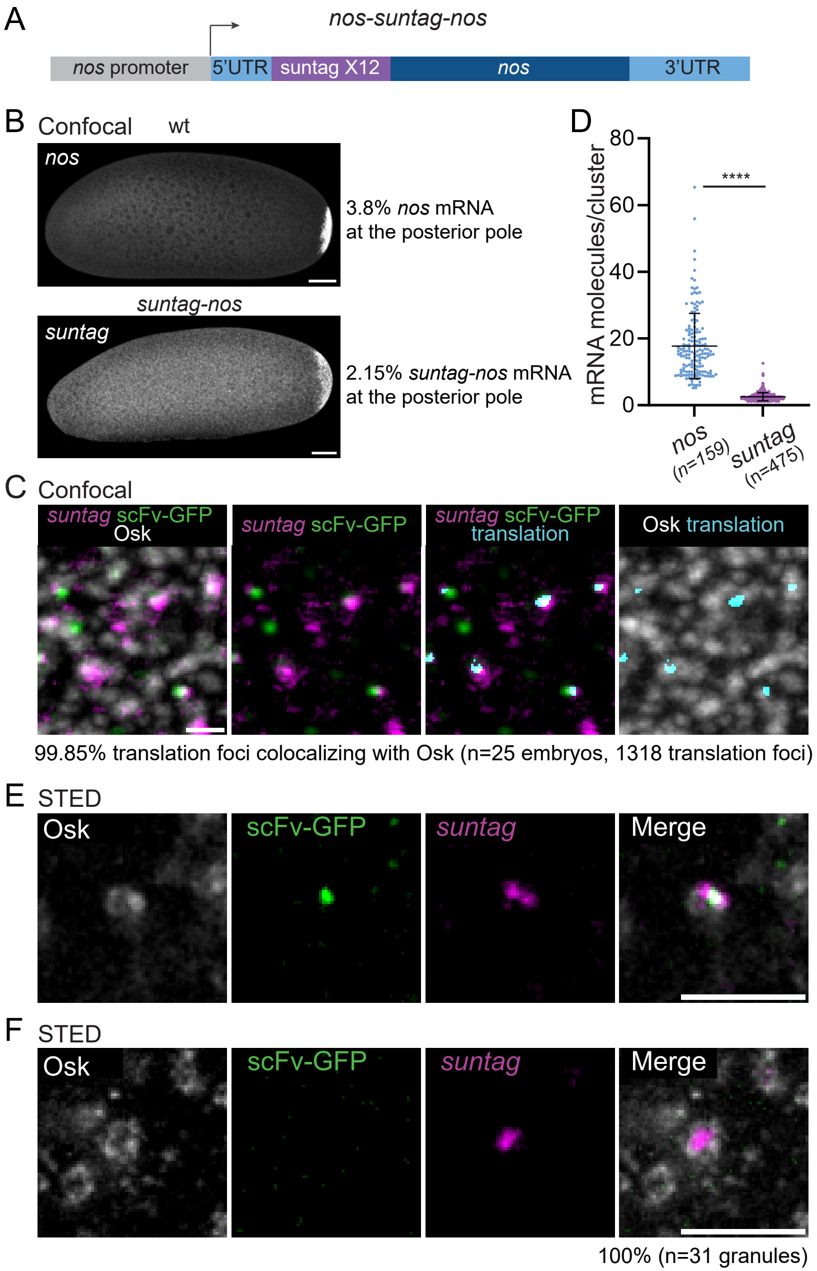

A
nos-suntag-nos
nos promoter
5'UTR
suntag X12
nos
3'UTR
B
Confocal
wt
nos
3.8% nos mRNA
at the posterior pole
suntag-nos
suntag
2.15% suntag-nos mRNA
at the posterior pole
D
mRNA molecules/cluster
80
60
40
20
0
****
nos
(n=159)
suntag
(n=475)
C
Confocal
suntag scFv-GFP
Osk
suntag scFv-GFP
suntag scFv-GFP
translation
Osk translation
99.85% translation foci colocalizing with Osk (n=25 embryos, 1318 translation foci)
E
STED
Osk
scFv-GFP
suntag
Merge
F
STED
Osk
scFv-GFP
suntag
Merge
100% (n=31 granules)


Figure S3

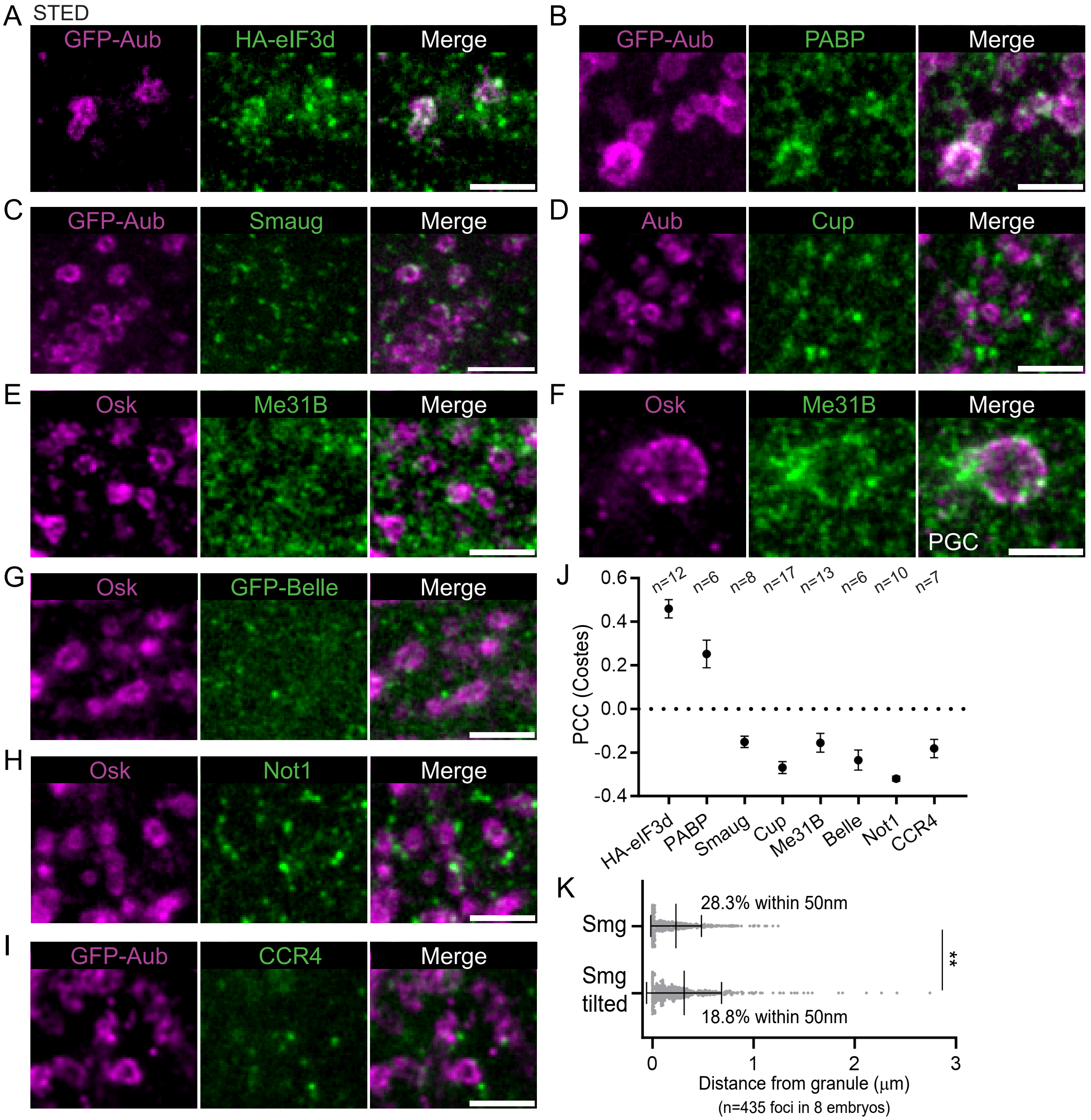

A
STED
GFP-Aub
HA-eIF3d
Merge
B
GFP-Aub
PABP
Merge
C
GFP-Aub
Smaug
Merge
D
Aub
Cup
Merge
E
Osk
Me31B
Merge
F
Osk
Me31B
Merge
PGC
G
Osk
GFP-Belle
Merge
H
Osk
Not1
Merge
I
GFP-Aub
CCR4
Merge
J
PCC (Costes)
0.6
0.4
0.2
0.0
-0.2
-0.4
n=12
n=6
n=8
n=17
n=13
n=6
n=10
n=7
HA-eIF3d
PABP
Smaug
Cup
Me31B
Belle
Not1
CCR4
K
Smg
Smg tilted
28.3% within 50nm
18.8% within 50nm
**
0
1
2
3
Distance from granule (μm)
(n=435 foci in 8 embryos)


Figure S4

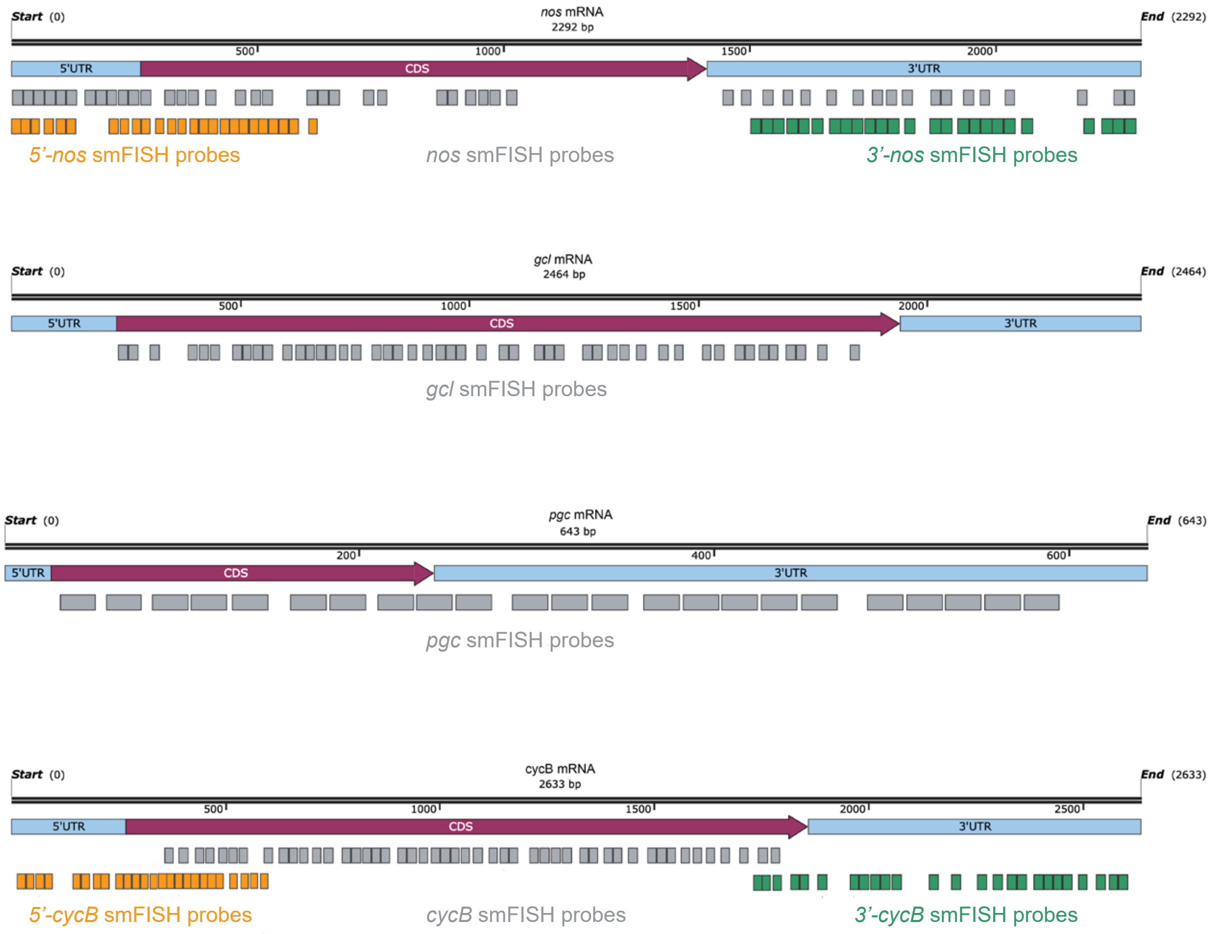

Start (0)
nos mRNA
2292 bp
End (2292)
500
1000
1500
2000
5'UTR
CDS
3'UTR
5'-nos smFISH probes
nos smFISH probes
3'-nos smFISH probes
Start (0)
gcl mRNA
2464 bp
End (2464)
500
1000
1500
2000
5'UTR
CDS
3'UTR
gcl smFISH probes
Start (0)
pgc mRNA
643 bp
End (643)
200
400
600
5'UTR
CDS
3'UTR
pgc smFISH probes
Start (0)
cycB mRNA
2633 bp
End (2633)
500
1000
1500
2000
2500
5'UTR
CDS
3'UTR
5'-cycB smFISH probes
cycB smFISH probes
3'-cycB smFISH probes


Figure S5

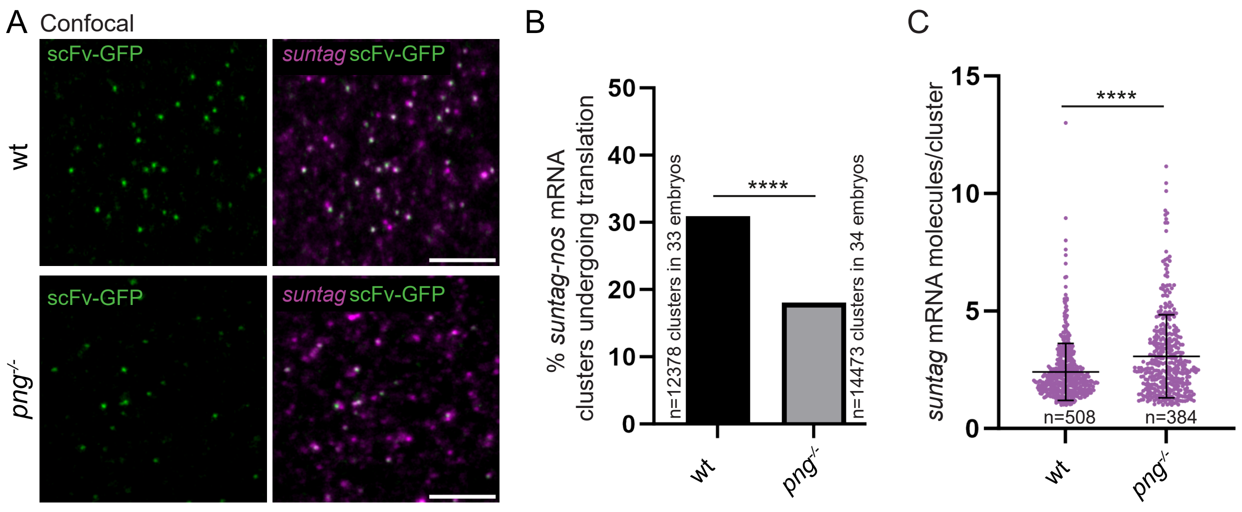
A
Confocal
scFv-GFP
suntag scFv-GFP
wt
png-/-
B
% suntag-nos mRNA clusters undergoing translation
n=12378 clusters in 33 embryos
n=14473 clusters in 34 embryos
****
0
10
20
30
40
50
wt
png-/-
C
suntag mRNA molecules/cluster
****
0
5
10
15
n=508
n=384
wt
png-/-

Figure S6

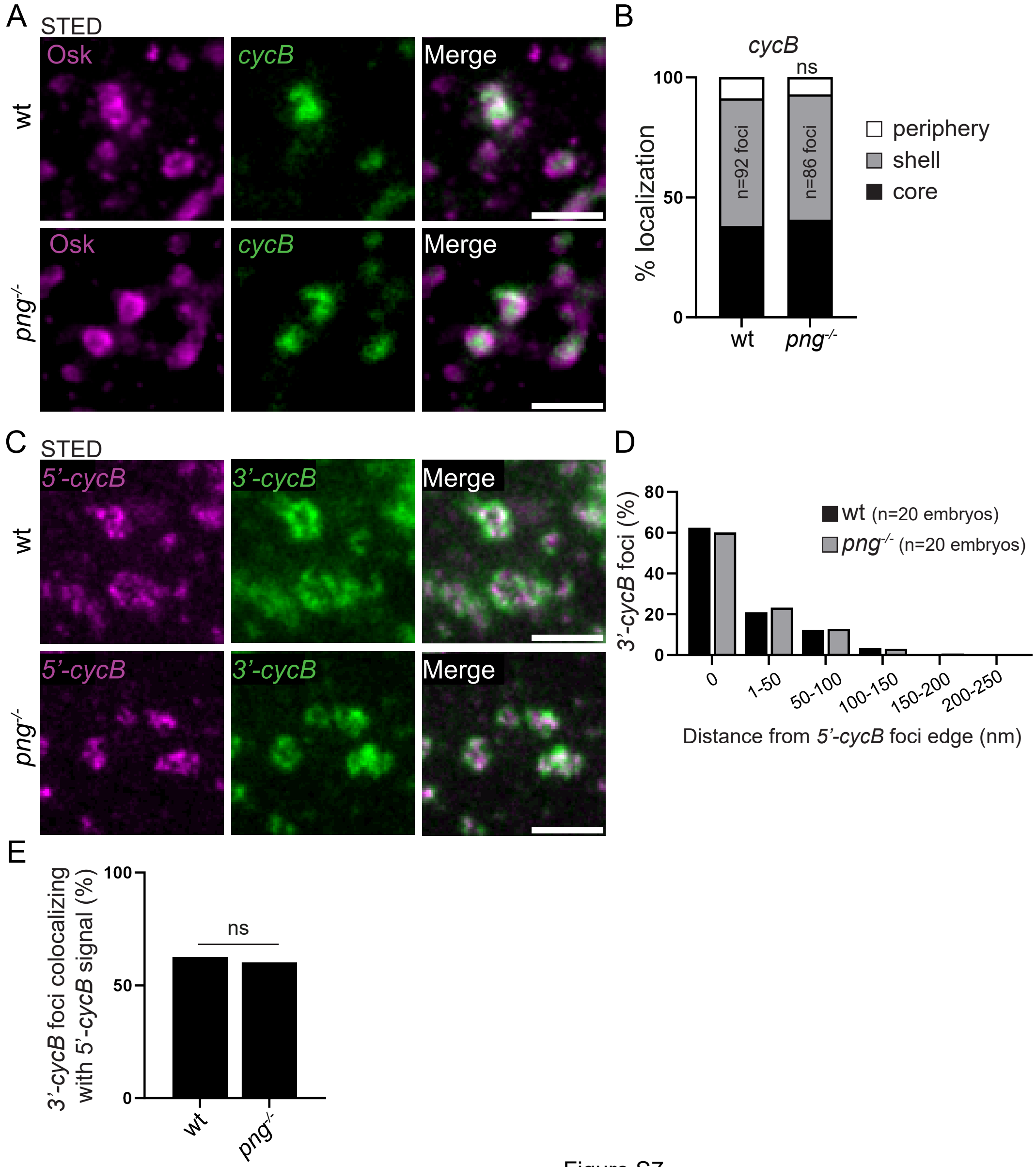
A
STED
Osk
cycB
Merge
wt
png-/-
B
cycB
ns
100
50
0
% localization
n=92 foci
n=86 foci
wt
png-/-
periphery
shell
core
C
STED
5'-cycB
3'-cycB
Merge
wt
png-/-
D
80
60
40
20
0
3'-cycB foci (%)
wt (n=20 embryos)
png-/- (n=20 embryos)
0
1-50
50-100
100-150
150-200
200-250
Distance from 5'-cycB foci edge (nm)
E
100
50
0
3'-cycB foci colocalizing with 5'-cycB signal (%)
ns
wt
png-/-

Figure S7

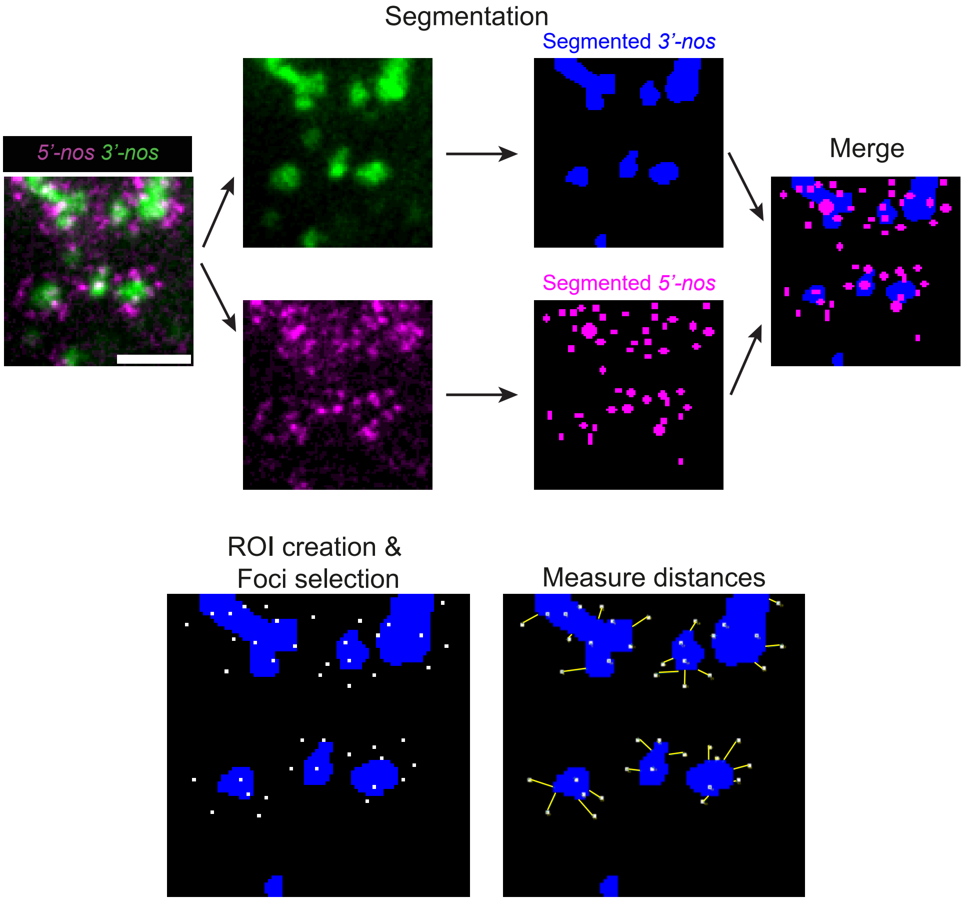
Segmentation
Segmented 3'-nos
5'-nos 3'-nos
Merge
Segmented 5'-nos
ROI creation &
Foci selection
Measure distances

Figure S8

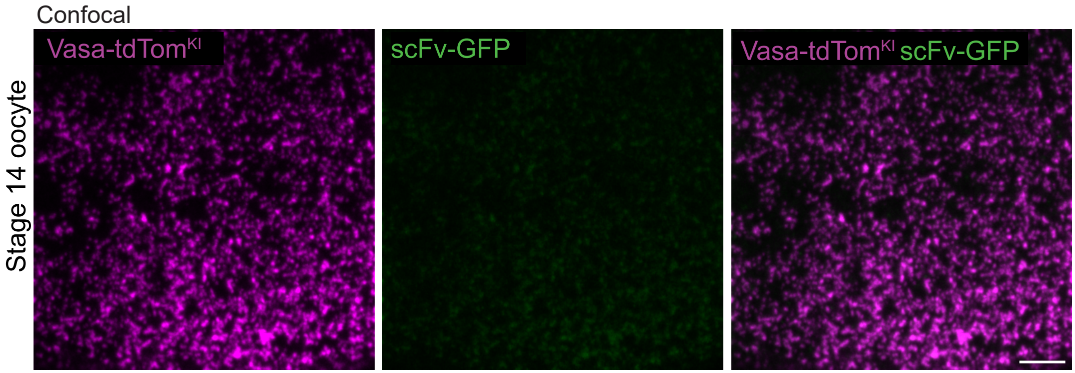
Confocal
Stage 14 oocyte
Vasa-tdTom$^{KI}$
scFv-GFP
Vasa-tdTom$^{KI}$ scFv-GFP

Figure S9

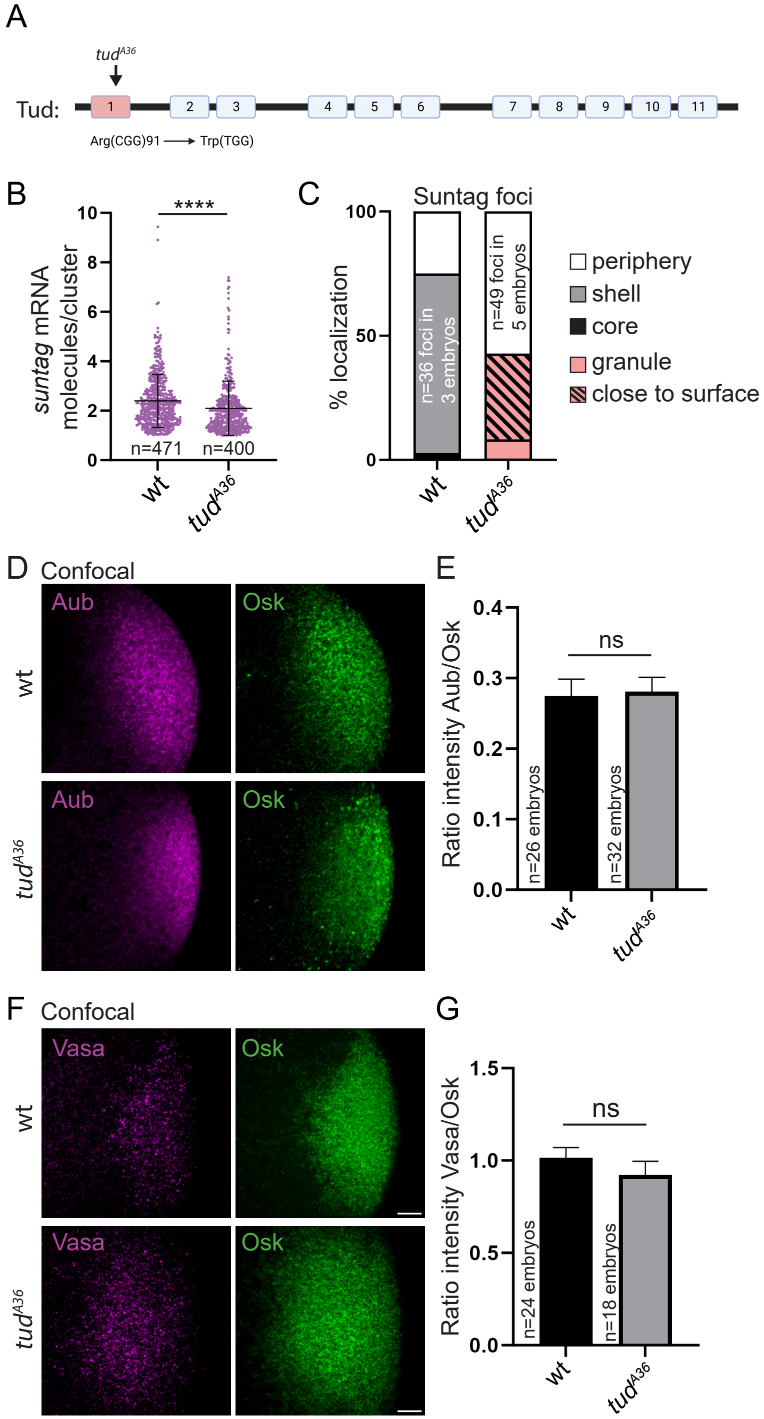
A
tudA36
Tud:
1
2
3
4
5
6
7
8
9
10
11
Arg(CGG)91 → Trp(TGG)
B
suntag mRNA molecules/cluster
****
n=471
n=400
wt
tudA36
C
Suntag foci
% localization
n=36 foci in 3 embryos
n=49 foci in 5 embryos
periphery
shell
core
granule
close to surface
wt
tudA36
D
Confocal
Aub
Osk
wt
tudA36
E
Ratio intensity Aub/Osk
ns
n=26 embryos
n=32 embryos
wt
tudA36
F
Confocal
Vasa
Osk
wt
tudA36
G
Ratio intensity Vasa/Osk
ns
n=24 embryos
n=18 embryos
wt
tudA36

Figure S10
